# Ethology of morphogenesis reveals the design principles of cnidarian size and shape development

**DOI:** 10.1101/2021.08.19.456976

**Authors:** Anniek Stokkermans, Aditi Chakrabarti, Ling Wang, Prachiti Moghe, Kaushikaram Subramanian, Petrus Steenbergen, Gregor Mönke, Takashi Hiiragi, Robert Prevedel, L. Mahadevan, Aissam Ikmi

## Abstract

During development, organisms interact with their natural habitats while undergoing morphological changes, yet it remains unclear whether the interplay between developing systems and their environments impacts animal morphogenesis. Here, we use the cnidarian *Nematostella vectensis* as a developmental model to uncover a mechanistic link between organism size, shape and behavior. Using quantitative live imaging, including extensive behavioral profiling, combined with molecular and biophysical experiments, we demonstrate that the muscular hydraulic machinery that controls body movement directly drives larva-polyp morphogenesis. Unexpectedly, size and shape development are differentially controlled by antagonistic muscles. A simple theoretical model shows how a combination of slow-priming and fast-pumping pressures generated by muscular hydraulics acts as a global mechanical regulator that coordinates tissue remodeling. Altogether, our findings illuminate how dynamic behavioral modes in the environment can be harnessed to drive morphogenetic trajectories, establishing ethology as a critical component of organismal morphogenesis – termed ethology of morphogenesis.

## Introduction

Developing metazoans generally confront two distinct environments (Gilbert and Epel, 2015). In early life, embryos are usually self-contained and develop in confined spaces such as gelatinous substances, eggshells, or parental hosts. Once they emerge from these protective compartments, developing animals transition into open environmental spaces. Here, they typically continue morphogenetic processes that vary from growth to metamorphic transformation (Flatt and Heyland, 2011) while simultaneously engaging in an extensive repertoire of organismal behaviors (Bateson, 1981; Pujala and Koyama, 2019; West et al., 2003). These behaviors include escaping from predators, searching for food, and finding a suitable ecological niche, all of which depend on animal motility. Although changes in behavioral patterns are integral to the early life history of metazoans, it remains unknown whether behavior in the open environment directly impacts morphogenesis (Bertossa, 2011; Prakash et al., 2021).

Thus far, the majority of leading morphogenesis studies have focused on providing in-depth, high spatial resolution views of early development, yielding insights into how embryonic mechanisms drive animal development at the single-cell resolution and a wide range of time scales (Collinet and Lecuit, 2021; Heisenberg and Bellaïche, 2013; Krzic et al., 2012; Stooke-Vaughan and Campàs, 2018; Yue et al., 2020). In contrast, behavioral studies commonly track whole-organism dynamics in terms of body actions in the environment over coarser time scales (Branson et al., 2009; Mearns et al., 2020). These differences preclude following the links between behavior and whole-organism coordination of morphogenesis in a freely developing animal, and reflect limitations of technologies and model systems used in development and behavior research. To study the role of behavior in development, an ideal system is one that can be imaged with the resolution typical for morphogenetic studies, manipulated genetically, and amenable to tracking its behavioral dynamics over long times.

The sea anemone *Nematostella vectensis* is a well-suited model system to examine the functional relationships between morphogenesis and behavior, given its relatively simple body plan, genetic tractability and optical accessibility (He et al., 2018; Ikmi et al., 2014; Layden et al., 2016; Renfer and Technau, 2017). *Nematostella* is composed of two thin tissue layers surrounding a single fluid-filled cavity with an oral opening (Layden et al., 2016; Martindale et al., 2004; Steinmetz et al., 2017), and undergoes larva-polyp transformation in an open aquatic space (Hand and Uhlinger, 1992). Previous work has shown that developing larvae engage in axial elongation that progressively transforms the initial ovoid morphology into a tubular polyp, while concomitantly exhibiting discrete behaviors such as cilia-based swimming, settlement, and muscle-driven body movements (Hand and Uhlinger, 1992) (Figure 1A), similar to other cnidarian species (Jackson et al., 2002; Müller and Leitz, 2002). While the inductive signals of larva-polyp transformation are an active area of research (Grasso et al., 2011; Levy et al., 2021a; Strader et al., 2018), and a growing number of whole-organism cell atlases are available for Cnidaria (Levy et al., 2021b; Sebé-Pedrós et al., 2018; Siebert et al., 2019), a predictive understanding of the cell-regulatory network that controls organismal morphogenesis is still missing.

**Figure 1.**
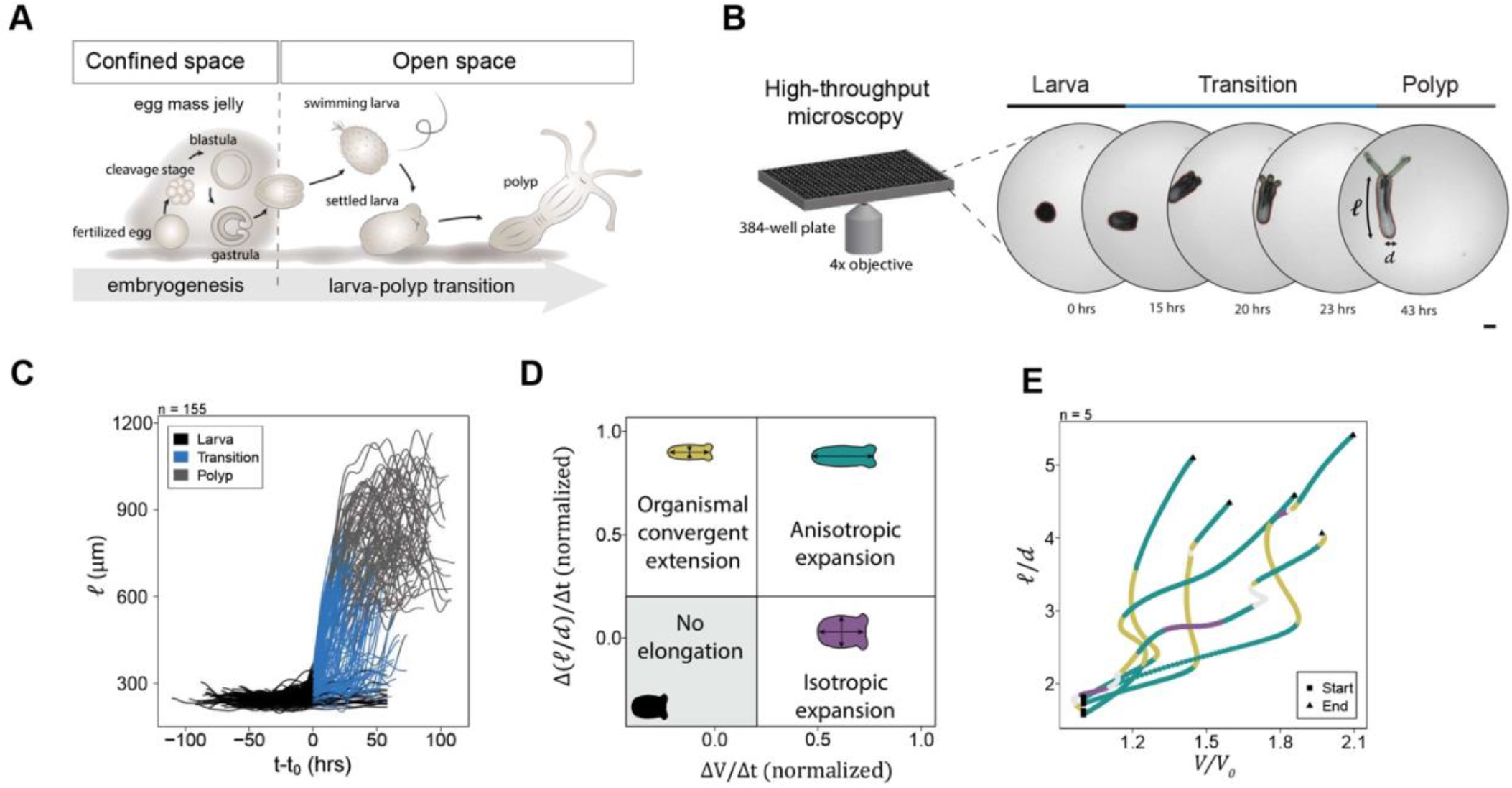
High-throughput live imaging reveals variability in morphodynamics during larva-polyp transition. (A) Schematic of *Nematostella* development showing an eventual transition from a motile to a sessile form that transforms from a spheroidal larva to an elongated polyp form. (B) Experimental setup and illustrative images for high-throughput recording of developing animals. Animal length (*ℓ(t)*), and diameter (*d(t)*) are functions of time and are used to compute animal aspect ratio. (C) Smoothed curves for body column length (*ℓ(t)*), shifted in time by t_0_, which marks the start of larva-polyp transition (*n = 155* animals). (D) Schematic showing definition of distinct morphodynamics, based on normalized volume and aspect ratio change. (E) Smoothed larva-polyp transition trajectories in relative size-shape morphospace for five animals. Colors indicate morphodynamics. Scale bar: 100 μm.

Here we propose ethology of morphogenesis, a concept that addresses how behavior and morphology development impact each other. To do so, we implemented advanced, custom-made optical imaging strategies, combined with computational, biophysical and molecular approaches, to map connections between size and shape development during axial elongation and behavioral changes that accompany larva-polyp transition. We show that the very same muscular hydraulic machinery that controls body movement in the environment drives system-level morphogenesis. We also introduced a simple mathematical model to quantify the role of the static-priming and dynamic-pumping hydraulic pressure in driving larva-polyp morphogenesis. Complementing this model, physical experiments with reinforced balloons qualitatively capture the phase space of organismal size and shape resulting from the perturbation of distinct muscle types. Therefore, our work establishes a direct mechanistic link between animal ethology and morphogenesis at both cellular and biophysical levels, and suggests that active muscular hydraulics plays a broad role in the design principle of soft-bodied animals.

## Results

### Larva-polyp transition shows variability in morphodynamics

To monitor the morphogenesis of freely developing animals at the population level and build correlation maps of morphogenetic and ethological states, we established a high-throughput live imaging method that captures larva-polyp transformation in a 384 well plate (Video S1; Figure 1B, n = 669 animals). As the development of *Nematostella* is asynchronous, we used changes in organismal circularity (a perfect circle is 1) as a geometrical feature to classify the developmental stages into larva (circularity <0.8), larva-polyp transition (circularity of 0.3-0.8), or polyp (circularity >0.3) (Figures 1C, and S1A and S1B). By tracking circularity over time, we observed that it decreases dramatically during larva-polyp transition, due to a three- to four-fold change in the length of the body and the development of oral tentacles (Figures 1C and S1A and S1B). At the population scale, the dynamics of larva-polyp transition were highly variable and the overall time for development ranged from 16 to 30 hours in most cases (Figures 1C and S1C). This temporal variability was associated with substantial changes in the elongation dynamics of individual organisms (Figure S1D) as well as different elongation rates across animals, and was independent of the disparity in body lengths (Figure S1E).

In order to quantify the dynamics of elongation, which links body size and shape, we developed a computational pipeline that maps the morphodynamics of live animals (see Methods and Figure S2). Our quantitative analysis showed that transforming larvae experience at least three morphodynamic events: 1) isotropic expansion due to body volume change that leads to an increase in overall size, 2) organismal convergent-extension (OCE) that leads to changes in shape, i.e. increase in body aspect ratio, and 3) anisotropic expansion by simultaneously increasing both body volume and aspect ratio (Figure 1D). While organismal convergent-extension and anisotropic expansion were the dominant developmental patterns observed, their temporal sequence and duration varied across developing animals (Figure 1E). Together these data suggest that the larva-polyp transformation is guided by a relatively plastic developmental program.

### Coordination of larva-polyp morphogenesis with behavioral modes

To examine behavioral changes and link them to observed morphodynamics, we tracked the patterns of settlement associated with switching between motile and sessile behaviors (Figures 2A and S3). In marine invertebrate larvae, this behavior marks a shift from a pelagic to a benthic form that can adhere to a substrate (Jackson et al., 2002; Leitz, 1997). We found that larvae can engage substrate in two states: a rapidly elongating mode wherein the larva is relatively sessile, and a slowly elongating state wherein the larva is highly motile (Video S2 and S3; Figures 2A and 2B, and S3). Rapidly elongating animals showed a clear period of continuous attachment to the substrate at the aboral region while the oral pole was free (Figure 2A and 2B). Interestingly, rapidly elongating larvae often exhibited a stereotyped morpho-ethological signature composed of anisotropic expansion, which is associated with a muscle-driven pumping behavior (Video S4; Figure 2C), followed by organismal convergent extension with a marked phase of sessile adhesion (Video S2; Figure 2D). This elongation pattern was tempered in settled larvae showing high motility (Figure 2C). Together, these results show a robust correlation of morphodynamics with behavioral patterns during larva-polyp transformation.

**Figure 2.**
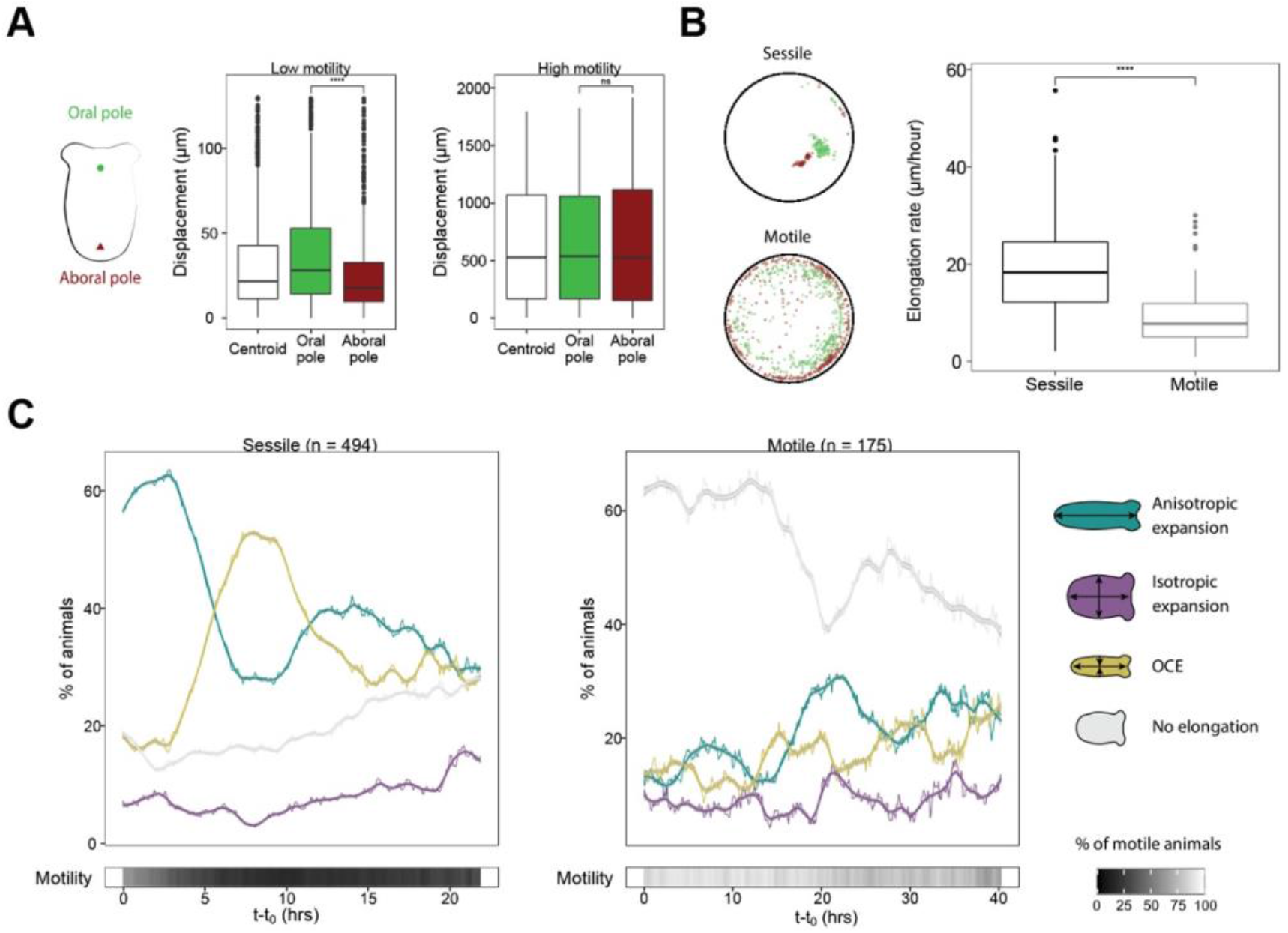
The pattern of morphodynamics correlates with behavioral modes during larva-polyp transition. (A) Measurements of oral and aboral pole displacement during low and high motility (two-sided unpaired Wilcoxon rank sum test; ****p < 0.0001). (B) Left: projection of oral and aboral pole location in a sessile and motile animal during the transition stage. Right: average axial elongation rate in sessile and motile animals (two-sided unpaired Wilcoxon test; ****p < 0.0001). (C) Time-shifted distribution of morphodynamics for sessile and motile animals (*n = 494* sessile animals, *n* = 175 motile animals).

Cnidarians have the morphology of a thin fluid-filled shell, and hydraulic forces due to luminal pressure are known to drive a variety of polyp behaviors involving support and movement (Batham and Pantin, 1950a; Kier, 2012). When the body wall has a uniform thickness and rheology, and no active contractility, changes in the intra-luminal pressure will only change organismal size. However, by changing the static properties of the wall, e.g., by modulating its thickness, or its dynamics via muscular activity, the organism can also change its shape. Indeed, the hydrostatic skeleton of cnidarian polyps consists of a muscular body wall that transmits muscular forces via dynamical fluid pressurization of the luminal cavity (Batham and Pantin, 1950a, 1951; Jahnel et al., 2014; Kier, 2012). The geometry of the body then serves to modulate the global pressure and leads to local reversible deformations of the polyp (Batham and Pantin, 1950a, 1951; Kier, 2012). This suggests that changing body wall tonicity in primary polyps should change organism morphology. To test this, we used linalool (Goel et al., 2019) to induce muscle relaxation in primary polyps, and observed that the body shape reverted to a more spherical morphology (Video S5), which demonstrates that static basal anisotropic muscular tonicity of the wall is required for maintaining the tubular shape. Given that transient muscular forces associated with body movement act in addition to this basal tonicity, and are correlated with the larva-polyp transition, we hypothesized that there is a causal relation between behavioral dynamics of muscular hydraulics and the morphological transition.

### Muscular-hydraulics drives larva-polyp morphogenesis

Prior to settlement, *Nematostella* larvae already possess a primordial hydrostatic skeleton that further expands its luminal cavity during larva-polyp transition (Jahnel et al., 2014). To measure the relative contributions of tissue and luminal volume increase to body size, we acquired 3D morphological data in live larvae and their corresponding polyps using custom-made dual-view optical coherence microscope (Figure S4A). The strong, label-free contrast between the tissue and the water-filled cavity permitted quantitative 3D mapping at a high and near-isotropic spatial resolution, even in more light scattering larval stages, due to the dual-view geometry (see Methods). We observed that although total tissue volume changed only around 1.5-fold, the body cavity volume increased approximately 9-fold (Figure 3A). We noted that fluid uptake through inflation of the body cavity is a major contributor to final polyp size; however, the uptake varies from one animal to another (Figure S4B). Overall, the fluid uptake causes an increased luminal pressure that leads to quasi-static body wall stresses.

**Figure 3.**
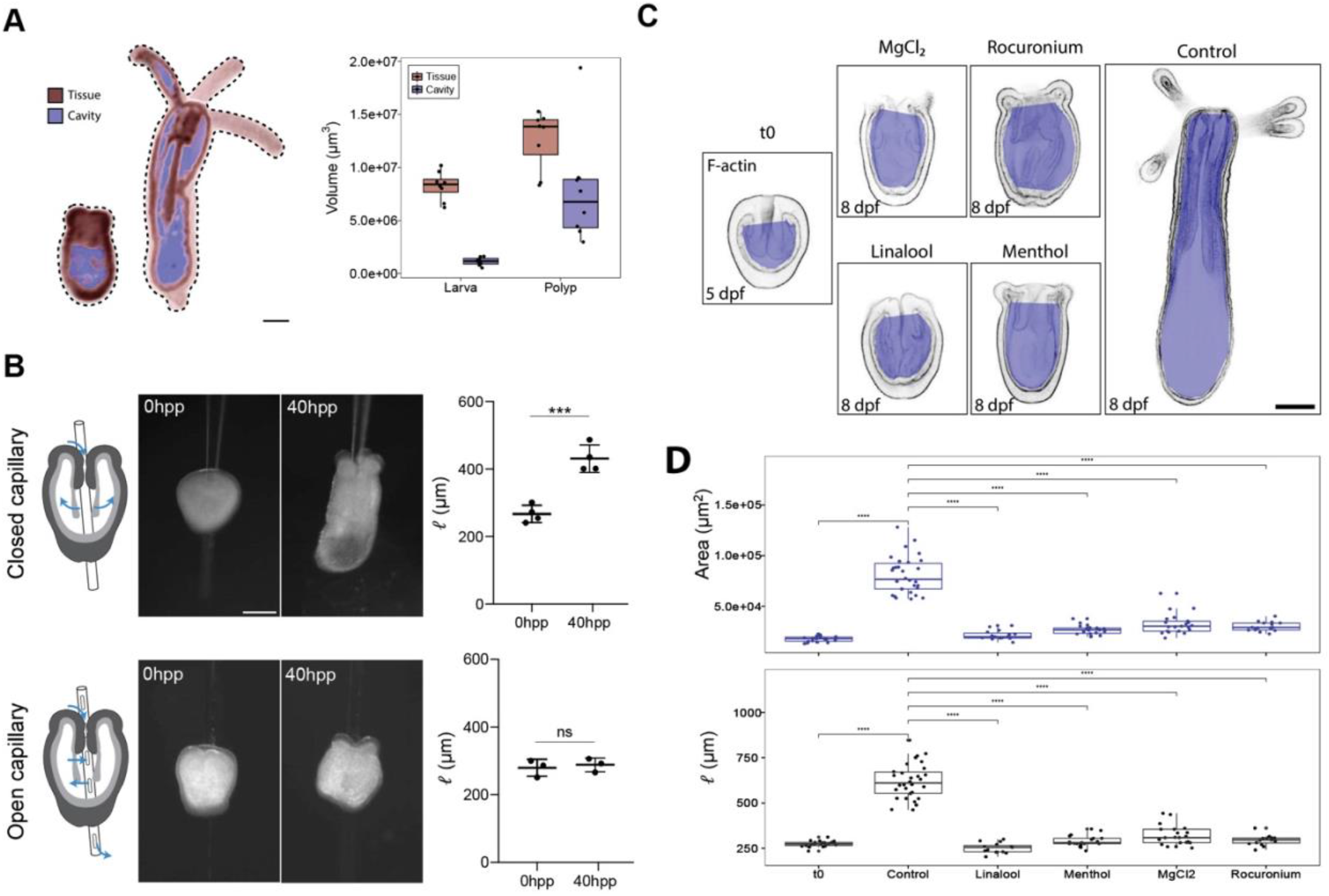
Muscular-hydraulics is required for cavity inflation and body elongation. (A)Left, mid-plane sections of 3D views of OCM images of the same animal at larva and polyp stage, segmented into tissue (red) and cavity (blue). Right: quantification of tissue and cavity volume at larva and polyp stage (n = 8). (B) Mechanical perturbation of cavity pressure by insertion of a closed capillary (control), or an open capillary, preventing build-up of pressure (two-tailed unpaired t-test, ***p<0.001; n.s p>0.05). hpp: hours post perforation. (C) Mid-plane sections showing animals stained for F-actin with phalloidin and fixed at 5 dpf (t0) or 8 dpf (drug-treated and control). Blue color indicates the area of the cavity plus internal tissue, as quantified in D. (D) Quantifications of body length (left) and cavity area in the mid-plane section (right) of drug-treated animals and controls (two-sided unpaired Wilcoxon rank sum test; ****p < 0.0001). t0: *n = 20*, control: *n = 30*, 1 mM linalool: *n = 15*, 400 μM menthol: *n = 19,* 0.5% MgCl_2_: *n = 21*, 0.5mM rocuronium bromide: *n = 15.* Scale bars: 100 μm.

To examine the role of hydraulic stresses during development, we inserted a hollow glass capillary with or without holes along its side through the mouth and the body column (Figure 3B). Capillaries with holes were used to facilitate water exchange between cavity and the external environment, and thereby prevent buildup of pressure inside the cavity. Control animals pierced with a closed capillary successfully underwent larva-polyp transformation. In contrast, larvae punctured with an open capillary failed to elongate, arresting their development at the budded stage. In parallel, we pharmacologically inhibited muscle function in settled larvae with different muscle relaxants (Abrams et al., 2015; Batham and Pantin, 1950a; Goel et al., 2019). Most drug-treated animals developed tentacle buds, but neither elongated nor inflated the body cavity (Figure 3C and 3D), thus mimicking the developmental phenotype caused by disruption of hydraulic stresses (Figure 3B). Taken together, these results show a crucial role of muscular-hydraulics in larva-polyp transformation.

### The muscular system differentially controls size and shape development

The muscle system of *Nematostella* consists of azimuthally oriented circular muscles, and axially oriented longitudinal parietal and retractor muscles (Jahnel et al., 2014) (Figure 4A and 4B). These muscles act antagonistically with respect to each other: contraction of circular muscles reduces the diameter of the body column, thereby forcing its length to increase in order to accommodate the same amount of fluid. In contrast, longitudinal muscle contraction reduces the length of the body column, forcing the widening of its diameter (Batham and Pantin, 1950a; Kier, 2012). As body contractions are muscle orientation-dependent, we investigated the contribution of different muscle types to morphogenesis using a shRNA knockdown (KD) screen targeting transcription factors associated with muscle development (Cole et al., 2020; Steinmetz et al., 2017). We identified that the orthogonal organization of the parietal and circular epitheliomuscular cells is severely disrupted in the KD of two paralogs of the Tbx20 family (*NvTbx20.1* and *NvTbx20.2*) (Figure 4C). As a result, the Tbx20 KD animals lacked a clear cylindrical shape and instead adopted a bulky sac-like polyp morphology, in which the aboral region from the mid-body was often most severely affected (Figure 4C). During development, the Tbx20 KD larvae changed their morphodynamics towards isotropic and anisotropic expansion, which increased organismal size through cavity inflation whereas aspect ratio increase was limited (Video S6; Figures 4D and S5). We independently validated the developmental function of the Tbx20 family using CRISPR/Cas9-mediated genome editing. Interestingly, F0 Tbx20 mutant animals exhibited a nearly spherical morphology resulting from a global disruption of parietal and circular epitheliomuscular cells (Figure S6A and S6B). In addition, spatially restricted muscle defects in Tbx20 mosaic mutant animals were sufficient to induce bleb-like deformations in the body wall (Figure S6A and S6B). These experiments demonstrate key roles for the Tbx20 family in controlling parietal and circular muscle patterning, and reveal a previously unknown link between the spatial organization of epitheliomuscular cells and organismal shape development.

**Figure 4.**
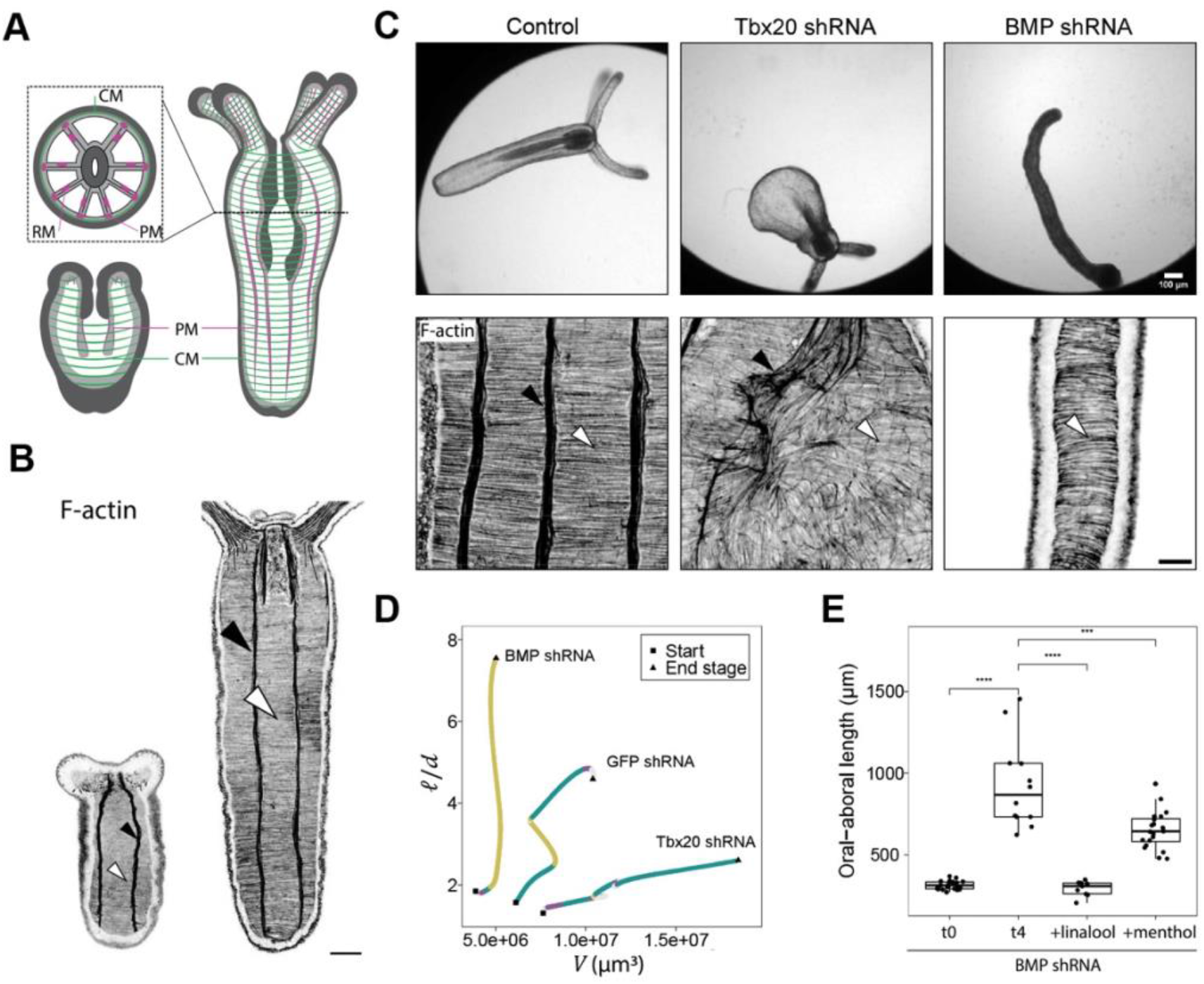
Antagonistic function of circular and longitudinal muscles controls size and shape development. (A) Schematic representation showing muscle organization in circular muscles (green) and longitudinal parietal and retractor muscles (magenta) in larva (left) and polyp (right). Dotted rectangle shows the cross-sectional view. CM: circular muscle, PM: parietal muscle, RM: retractor muscle. (B) F-actin staining showing parietal (black arrowhead) and circular (white arrowhead) epitheliomuscular cell organization in a transforming larva and primary polyp. (C) Top: brightfield images of control, Tbx20 KD and BMP KD animals. Bottom: representative confocal images of F-actin staining showing muscle organization in control (ordered both longitudinally-parietally and azimuthally-circularly), Tbx20 KD (disordered), and BMP KD (ordered azimuthally) animals. (D) Trajectories in morphospace for a control, Tbx20 KD, and a BMP KD animal, color-coded for morphodymanics (yellow: organismal convergent extension, purple: isotropic expansion, green: anisotropic expansion, grey: no elongation). (E) Quantification of body length for animals injected with BMP shRNA, measured at 2 dpf (t0), and 6 dpf (t4, drug-treated animals and control) (two-side unpaired Wilcoxon rank sum test; ****p < 0.0001, ***p < 0.001). t0: *n = 23*, DMSO control: *n = 12,* 1 mM linalool: *n = 9,* 400 μM menthol: *n* = 20. Scale bars: 50 μm (B), 20 μm (C).

Simultaneously, we also performed a KD of BMP, which leads to a complete loss of the secondary body axis including the mesenteries and their associated longitudinal (parietal and retractor) muscles while maintaining circular (azimuthal) muscle organization (Leclère and Rentzsch, 2014) (Figure 4C). This caused a dramatic change in the shape of BMP KD animals resulting in an elongated noodle-like form (Figure 4C). Consistent with this morphology, organismal convergent extension was the dominant driver utilized by the BMP KD larvae (Video S7; Figures 4D and S5). Furthermore, this shape change was blocked when BMP KD larvae were treated with muscle relaxants (Figures 4E and S7), showing the critical role of circular muscles in organismal convergent extension driven elongation. As coordinated body wall deformations can generate fluid uptake via the oral opening due to suction (Batham and Pantin, 1950a; Vogel, 2007), the lack of cavity inflation in the BMP KD could be due to defective muscle-driven fluid dynamics. Supporting this hypothesis, pumping behaviors correlated with volume increase in transforming wild type larvae (Video S4), and the inhibition of muscle function blocked cavity inflation (Figure 3C and 3D). Taken together with earlier results, these findings suggest that coordinated muscle contractions are critical drivers of changes in organismal size and shape during development.

### Muscular-hydraulics impacts tissue remodeling

Next, we examined the relationship between muscular-hydraulics and the multicellular dynamics associated with tissue elongation in the body-wall (Figure 5A), which are distinguished by three motifs: oriented cell division, cell shape change and oriented cell rearrangements (Fritz et al., 2013). We found that blocking cell proliferation with hydroxyurea during larva-polyp transition has minimal effect on axial elongation dynamics (Figure S8). Thus, larval tissue remodeling occurs primarily by modulating cell shape and position; cell shape changes reduce body-wall tissue thickness and increase the entire animal’s surface (Fritz et al., 2013) (Figure 5B), while oriented cell rearrangements account for the directional expansion along the oral-aboral axis (Fritz et al., 2013), as visualized by tracking photoconverted Kaede-positive tissue stripes in the epidermis (Figure 5C).

**Figure 5.**
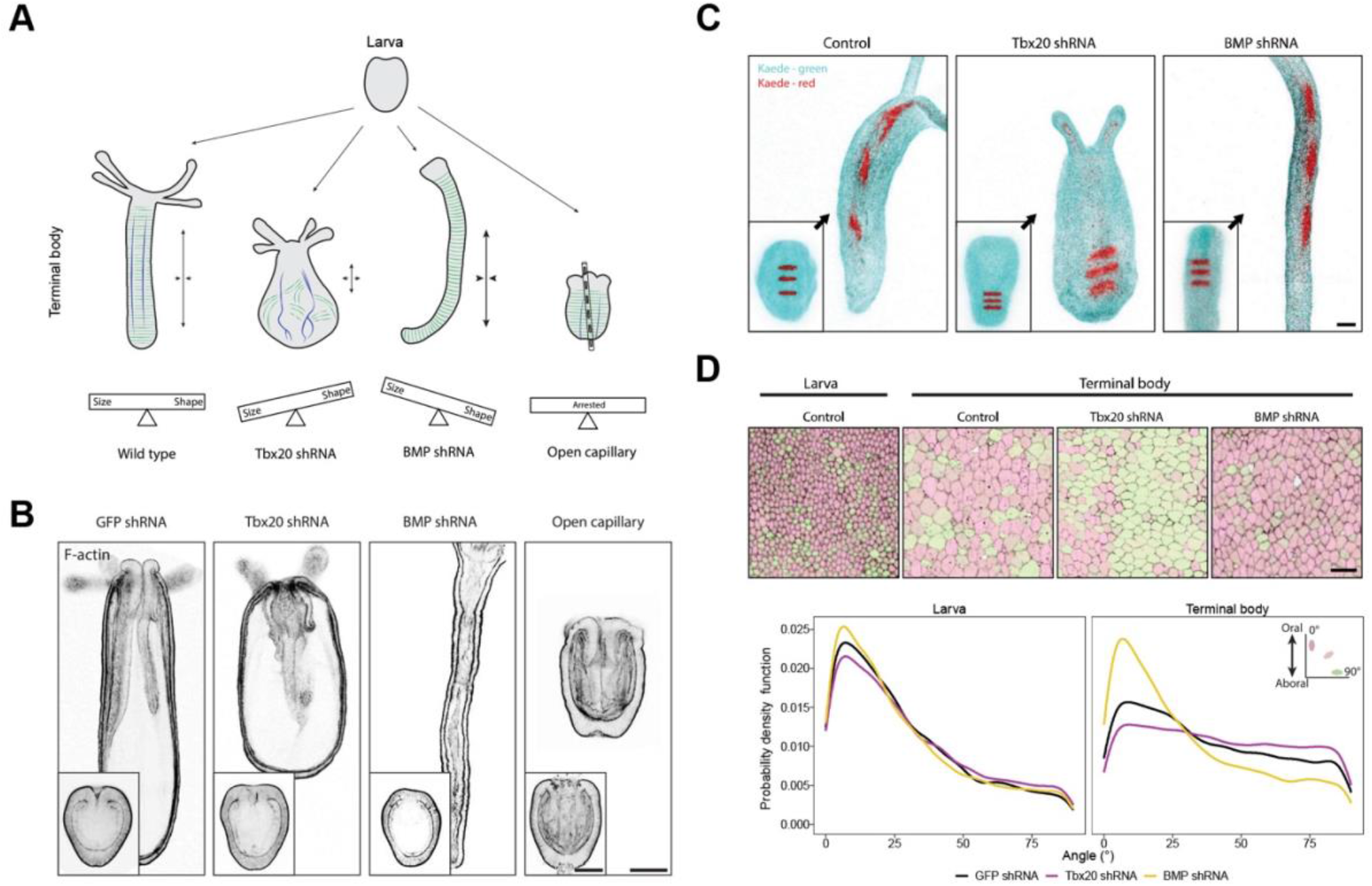
The cellular dynamics in the body wall depend on muscular-hydraulics. (A) Schematics of muscle organization and animal shape-size relationship for wild type, Tbx20 KD, BMP KD, and depressurized animals. (B) Mid-plane slices of animals stained for F-actin. Insets show larvae. (C) Photoconverted horizontal tissue stripes in larvae expressing Kaede (cyan: not converted; red: photo-converted), and their corresponding topological changes in developed wild type, Tbx20 KD, and BMP KD animals. (D) Quantification of cell orientation for larva and terminal body. Cell junctions were stained with an antibody for Cadherin3. Scale bars: 100 μm (B), 50 μm (C), 10 μm (D).

To determine how these morphogenetic changes drive tissue elongation, we analyzed the response of the epidermal cells in the body-wall to three distinct muscular-hydraulic stress conditions: 1) lack of hydraulic stress in depressurized animals; 2) axial stretching with minimal inflation in BMP KD animals; and 3) inflation with defective anisotropy in Tbx20 KD animals (Figure 5A). In all cases, larvae had indistinguishable epithelial thickness from controls, and most cell angles were oriented along the oral-aboral axis (Figure 5B and 5D). Reinforcing the importance of muscular hydraulics in development, physically deflated larvae maintained their epithelial architecture (Figures 5B and S9A), and their epidermal cells lacked apical actin-enriched rings that are typical to polyp cell differentiation (Figure S9B). In BMP KD animals, while larval cell orientation bias was maintained, we observed extensive oriented cell rearrangements along the main axis (Figures 5C and 5D, and S9C). In contrast, Tbx20 KD animals showed reduced cell rearrangements along the main axis and a broader distribution of cell angles (Figures 5C and 5D, and S9C). These patterns of tissue remodeling mirrored the anisotropic and isotropic stresses in BMP KD and Tbx20 KD animals, respectively. Therefore, these findings suggest a strong link between the developmental behavior of muscular hydraulics and the cellular morphogenetic responses of the body wall.

### Quantitative physical model of larva-polyp transformation

To quantitatively connect larva-polyp morphogenesis to the dynamic behaviors of muscular hydraulics, we introduced a minimal biophysical framework where we treat the larva as a thin elastic cylindrical shell, with a closed aboral end and an oral opening capable of ingesting fluid to increase cavity volume. For the rheological model of the wall, we assume a Kelvin-Voigt constitutive relation with an elastic element (Young’s modulus *E*) and a viscous dashpot (viscosity *η*) attached in parallel and a muscular spring (*E*_m_) attached in series with the *E*-*η* branch, leading to an elastic response at short time scales and slow creep at long time scales (Figure 6A). This is consistent with the observation that the tonicity of the muscles holds the shape of the polyps (Alexander, 1962) (Video S5). For a thin cylindrical shell with length *ℓ*(*t*), radius *r*(*t*), wall thickness *h*(*t*), the axial and azimuthal stresses are described by: σ_*Z*_ = *Pr*/2*h* and σ_*θ*_ = *Pr*/*h*, where *P*(*t*) is the intra-luminal pressure, which consists of a static component that is due to a basal muscle tone, fluid uptake, or osmotic imbalances, and a dynamic component due to muscular contractility. At the simplest level, we can write this as *P*(*t*) = *p*_*T*_ + *p*_0_ *Cos*(*ωt*) where *p*_*T*_ is the base pressure (static priming pressure) and *p*_0_ is the amplitude of the oscillatory pressure term with *ω* frequency (dynamic pumping pressure). Using a micropressure probe (Chan et al., 2019; Petrie et al., 2014), we measured the luminal pressure of primary polyps. The static luminal pressure was approximately 1,000 Pa, which increased to about 6,000 Pa in response to body contractions (Video S8; Figure 6C). When animals were treated with a muscle relaxant, this dynamic pressure increase was abolished (Video S9).

**Figure 6.**
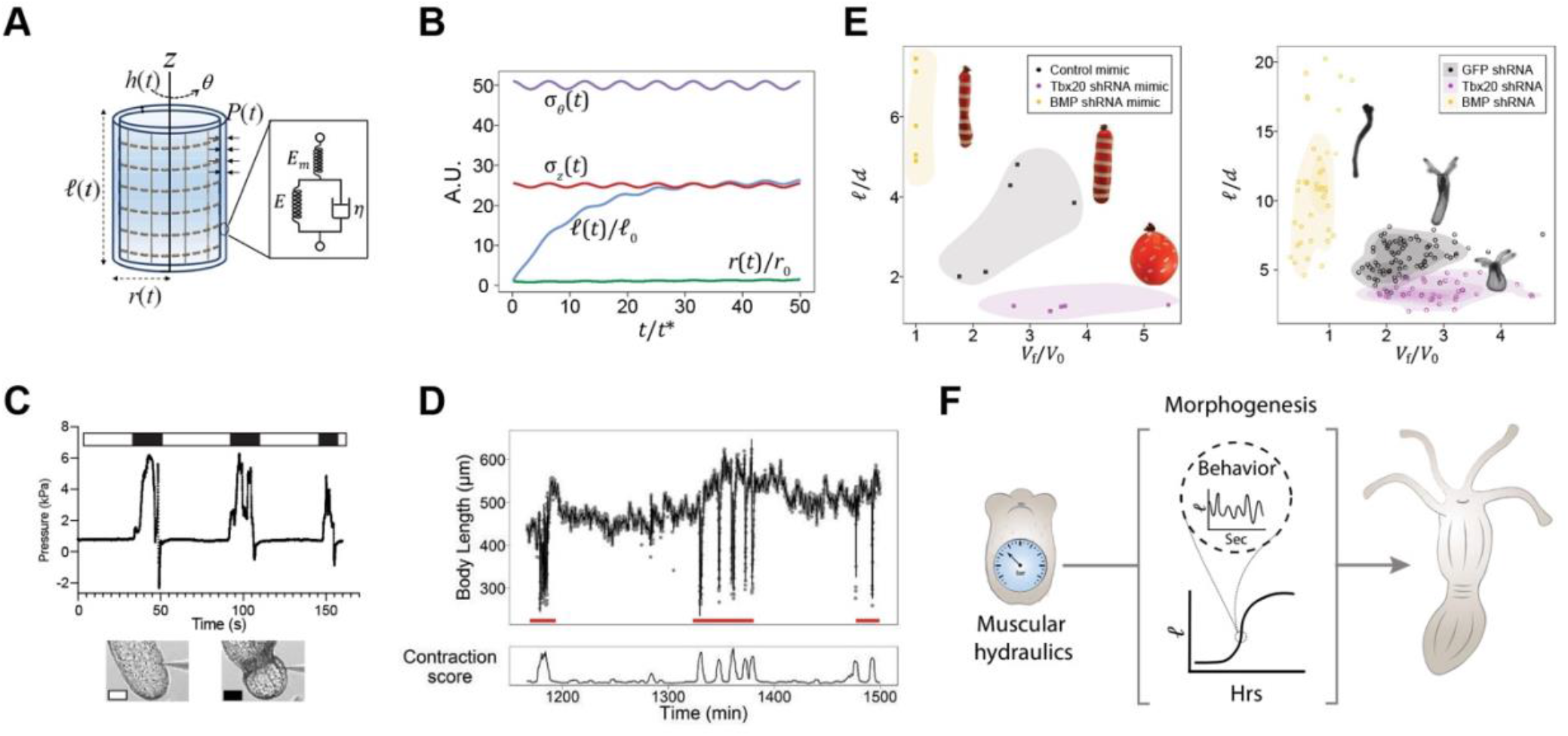
Biophysical model recapitulates larva-polyp size and shape transformation. (A) *Nematostella* modeled as a cylindrical shell of thickness *h(t)*, length *ℓ(t)* and radius *r(t)*, with time varying pressure in the cavity *P(t)*. A Kelvin Voigt model with a muscular spring in series is used to model tissue properties. (B) The resulting dynamic analytical model obtained by solving Eqs. A1–A4 (see Methods) to give the shape change quantified by *ℓ*(*t*)/*ℓ*_0_ and *r(t)*/*r*_0_, and the axial (σ_*z*_) and azimuthal (σ_*θ*_) stresses applied to the cylindrical shell as a function of dimensionless time (*t*/*t\**) at steady state, where *t*\*=*η*/*E*. (C) Cavity pressure in a contracting animal shows a characteristic pulsatile form with large variations. (D) Experimentally observed changes in axial length correlate with body contractions (red lines), consistent with a priming role for the average luminal pressure, and a pumping role for the dynamic muscular-hydraulic pressure. (E) Morphospace for body aspect ratio (*ℓ*/*d*) as a function of cavity volume changes (*V*_f_/*V*_0_) for balloon (left) and animal (right) experiments show similar qualitative trends for control, Tbx20 KD, and BMP KD conditions. Representative images of each condition are shown. (F) Summary of how muscular hydraulics drives the larva-polyp transformation.

In the limit of small strains and strain rates, we can use the Kelvin-Voigt linear constitutive relation to write down expressions for the axial and azimuthal strain and relate them to the corresponding stresses, as well as expressions for the cavity and tissue volume changes in terms of the geometry of the animal (see Methods). A qualitative view of the larva-polyp transformation is shown by solving the dynamic trajectories of body length *ℓ*(*t*)/*ℓ*_0_, radius *r*(*t*)/*r*_0_, and wall (tissue) thickness *h*(*t*)/*h*_0_ as a function of time, along with the principal stresses in the axial, σ_*z*_(*t*) and the azimuthal directions, σ_*θ*_(*t*) (Figure 6A and 6B). In this context, the static pressure brings the axial wall stress close to the yield stress (priming) and the dynamic oscillatory pressure term (pumping) tips it past the yield stress (Figure 6B), leading to an oscillatory elongation profile consistent with the correlation between axial length increase and body contractility (Figure 6D). Collectively, these results show that a combination of static “priming” luminal pressure and the dynamic “pumping” pressure causes the body wall to yield irreversibly, which directly links behavioral dynamics to larva-polyp morphogenesis.

### Reinforced balloons mimic the range of polyp shapes and sizes

Moving beyond the dynamics of body elongation towards the steady state shapes of the polyp, we note that under this limiting condition strain rates vanish (see Methods), and the wall tissue volume remains conserved. In Tbx20 KD animals, the disruption of parietal and circular muscles implies that their contribution towards the elastic stiffness becomes negligible, so that the axial and azimuthal stiffnesses become comparable. Combining these assumptions with the steady state equations of equilibrium yields the simple result that all strains are isotropic, i.e. Tbx20 KD larvae adopt a spherical polyp morphology that inflates isotropically (Figures 4C and S6). In contrast, BMP KD animals, the azimuthal muscles are conserved, while the axial muscles are lost, implying that effective stiffness in the azimuthal direction is much larger than the stiffness in the axial direction. This explains the origin of a thin and elongated morphology with a conserved cavity volume and large aspect ratio in BMP KD animals (Figures 4C and 5B). Finally, for wild type animals, while the body wall tissue volume remains conserved, the volume of cavity increases through fluid uptake. As a result, we obtain a relationship similar to that seen in the BMP KD phenotype, i.e., the aspect ratio can be larger than unity if the axial stiffness is lower than the azimuthal stiffness. To support the relationship between our minimal theoretical arguments and the described morphologies, we realized simple physical experiments with elastomeric balloons, with and without bands and tapes to mimic the passive effects of the constraining parietal (longitudinal) and circular (azimuthal) musculature. By varying the protocols for inflation and reinforcement, we reproduced the morphological patterns of Tbx20 KD, BMP KD and wild type animals using balloon mimics (Figure 6D and 6E). This further supports that our quantitative physical model, which incorporates static and dynamic pressure components, as well as directionality of the stress (axial vs. azimuthal), accurately recapitulates observed changes that accompany larva-polyp morphogenesis.

## Discussion

Understanding morphogenesis requires integrating molecular and cellular processes with large scale active deformations and flows at an organismal level. In particular, large scale hydraulic forces have recently been implicated in a number of early morphogenetic events in cellular cysts (Ruiz-Herrero et al., 2017), otic vesicles (Mosaliganti et al., 2019) and even whole embryos (Chan et al., 2019). In parallel, local muscle contractility is emerging as a key developmental player for many biological systems, including the gut (Huycke et al., 2019; Khalipina et al., 2019; Shyer et al., 2015), bone (Felsenthal and Zelzer, 2017; Schwartz et al., 2013), lung (Kim et al., 2015), whole embryos (Lardennois et al., 2019; Zhang et al., 2011) and wound healing and regeneration (Abrams et al., 2015; Livshits et al., 2017). By conceptualizing the ethology of morphogenesis as a key developmental parameter, here we show how the combined action of the muscular system and body hydraulics links animal behavior to the development of organism size and shape, and establish the critical role of muscular hydraulics in the larva-polyp transition.

During early development, the differentiation of a primordial hydrostatic skeleton consisting of a muscular body wall and a pressurized lumen produces a larva that behaves like a hydraulic pump (Figure 6F). Rapid contractions of the muscular wall in motile larvae generate reversible body deformations underlying discrete behaviors such as crawling, stretching and inflation/deflation. Over longer times, in settled larvae, muscular hydraulics form an active biomechanical module that leads to cavity inflation, which increases organism size. Local muscle organization and stresses in concert with the luminal pressure change body shape by imposing mechanical stresses that coordinate large scale tissue remodeling via slow luminal pressure priming and fast muscular hydraulic pumping. In many engineered systems, hydraulics is defined by the ability to harness pressure and flow into mechanical work, with long range effects in space-time. As animal multicellularity possibly evolved in an aquatic environment, we propose that early animals likely exploited the same physics, with hydraulics driving both developmental and behavioral decisions.

Our work also showed that a stable sessile adhesion at the aboral pole initiates a robust morpho-ethological pattern that drives axial elongation, suggesting that the variability in the elongation dynamics of larva-polyp transformation can be explained via the stochastic deployment of different behaviors during development. However, the nature of the morpho-etho coupling due to a combination of changes in fluid uptake rates, wall mechanical properties, or wall muscular activity remains an open question. The ubiquity of hydrostatic skeletons in the animal kingdom (Kier, 2012) and especially in marine invertebrates suggests a broad role for active muscular hydraulics on their developmental dynamics, and sets the stage for the study of the neuro-mechanical control of morphogenesis in organisms and mapping the behavioral dynamics and morphologies across species.

## Author contributions

A.S and A.I conceived the idea for this project and designed most of the experiments. A.C and L.M designed, generated and integrated the theoretical and physical models with the biophysical experiments. A.S established the high-throughput live imaging, and computational pipeline for image analyses with the support of G.M. A.S performed shRNA KDs, pharmacological treatments, staining, photoconversion, all imaging and data quantification. A.I performed the capillary experiments and CRISPR/Cas9 mutagenesis. P.M measured the cavity pressure under T.H’s supervision. L.W and R.P designed, built and optimized the dual-view OCM. K.S. acquired the OCM images, L.W, K.S and A.S performed image data processing, and A.S. segmented the images. A.C conducted the physical experiments with balloons. L.M. outlined the theoretical framework. P.S optimized the pharmacological experiments. A.I drafted the manuscript with inputs from A.S, A.C and L.M. All authors edited the manuscript.

## Acknowledgements

We thank C. Kresimir for animal husbandry, and the members of the ALMF at EMBL including B. Neumann, L. C. Schuetz, A. Halavatyi, S. Reither and S. Terjung for imaging support. We thank O. Matar for his help with shRNA microinjection, A. Paix for his support with genome editing, S. Cheung for generating the Kaede mRNA, and C.J. Chan for the discussions at the early stage of the work. We also thank C. Tischer and V. Uhlmann for their inputs in data analysis. We thank U. Technau for sharing the Cadherin 3 antibody. We also thank S. De Renzis, N. Petridou, G. Rapti, A. Erzberger, A. Ephrussi, and A. Aulehla for their comments on the manuscript. This work was supported by the NSF-Simons Center for Quantitative Biology at Harvard University and by the European Molecular Biology Laboratory.

## Declaration of interests

The authors declare no competing interests.

## Methods

### Animal husbandry

*Nematostella* adults were spawned every 3 weeks, and were maintained in a circulating system with 12 parts per thousand (ppt) artificial seawater (ASW) (sea salt; instant ocean) at 17 °C in the dark. Spawning was induced in a light box using a temperature of 28 °C and light intensity of 250–300 lumen per square foot for about approximately 6 hours (Genikhovich and Technau, 2009). Spawning occurred within 3–4?h after a cold-water change (17 °C).

### High-throughput live imaging

Larvae were placed in a 384 well plate (Corning, 3540) using a glass mouth pipet. The plate was imaged using an Acquifer screening microscope with a 4x magnification objective, with brightfield channel (20% intensity), at 5-minute time resolution at 27 °C for 3-7 days. For high-time resolution recordings, the time interval was 5 seconds.

### Image analysis of live animals

All image analyses were performed in FIJI (https://imagej.net/software/fiji/) (Schindelin et al., 2012). Development of live animals was analyzed using a Jython script written to run in FIJI. The script segments the animal and identifies the main body axis and secondary branches (tentacles), by using the AnalyzeSkeleton plugin (Arganda-Carreras et al., 2010). Body length is defined as the length of the skeleton branch that is identified as the main branch, plus a correction at the oral and aboral extremities based on the distance map of the segmented animal. Animal body column width is calculated from the distance from each point along the main body axis to the boundary of the animal using FIJI’s Euclidean distance map function (body column width = distance to boundary * 2). The volume is estimated by taking the sum of the volumes of 1-pixel tall cylinders with a diameter dependent on the local width, plus two half-spheres to account for the two poles. The script also makes use of the MorphoLibJ package (Legland et al., 2016) and the BioFormats plugin (Linkert et al., 2010). The windowed sinc filter from the pyBOAT package (Mönke et al., 2020), with cut-off periods of 500 and 1000 minutes was used to smooth the 5-minute time resolution data. This removes all high frequency components and leaves only the main trend intact. Staging of the animals was based on the decrease in circularity of the segmented 2D shape of the animal, which occurs as a result of elongation of the body column and outgrowth of tentacles. Circularity is defined as 4π(area/perimeter^2^), where a value of 1 corresponds to a perfect circle. A threshold of 0.8 for smoothed circularity, with cut-off period of 1000 minutes, was used to mark the boundary between larva and the larva-polyp transition. Similarly, a threshold of 0.3 marked the boundary between animals undergoing larva-polyp transition and polyps. For shRNA-injected animals, this staging method was refined to include body column volume and aspect ratio using the following thresholds: larva – transition: circularity: 0.8, aspect ratio: 1.8, body column volume: 0.75*10^7^ μm^3^; transition – polyp: circularity: 0.3. aspect ratio: 7, body column volume 1.8*10^7^ μm^3^.

To define different morphodynamics, the derivatives of the smoothed aspect ratio and estimated body column volume (cut-off period of 500 minutes) were used. The values were truncated and normalized to the 95^th^ percentile for aspect ratio and volume change. This way, all values are scored between −1 and 1, where 0 means no change and 1 corresponds to the value of the 95th percentile. Negative scores indicate a decrease in aspect ratio or volume. To define the different morphodynamics, a cut-off value of +0.2 was used, such that organismal convergent extension is defined by a normalized aspect ratio change ≥ 0.2 and body column volume change < 0.2, isotropic expansion is defined by normalized aspect ratio change < 0.2 and body column volume change ≥ 0.2, and anisotropic expansion is defined by normalized aspect ratio change and body column volume change ≥ 0.2. If none of these criteria are met, the morphodynamics at that given time point is labeled ‘no elongation’ to indicate that the animal either decreased or did not substantially increase in aspect ratio and/or volume.

To distinguish between high and low motility for a given animal at a given time point, we used a threshold at 130 μm displacement per 5-minute interval (Figure S3A). To distinguish sessile from motile animals, we used a threshold at 130 μm displacement for the median value of displacement during the entire transition stage.

To show contractions (Figure 6D), data was acquired at 5-second resolution and smoothed with a cut-off period of 40 seconds, using the sinc-smooth function (pyBOAT). As a proxy for contractility, the rolling sum (window of 250 sec) of the absolute change in smoothed circularity was used. This is based on the observation that the circularity of the animal’s 2D shape fluctuates as the animal contracts, hence the cumulative score in a given time window is predictive for the extent to which the animal deforms.

### Optical coherence microscopy and data processing

Optical Coherence Microscopy (OCM) has emerged as a promising modality for three-dimensional morphological imaging of small, behaving animals because of its high volumetric speed, label-free contrast and non-phototoxicity (Drexler and Fujimoto, 2015). Here we developed a bespoke OCM platform optimized for in-vivo imaging and biometry of non-anesthetized *Nematostella*. Because light scattering, absorption, and shadowing artefacts, especially at the larval stage, did not permit whole animal imaging, we engineered a tailored, dual-view spectral-domain Optical Coherence Microscopy (DV-OCM) system that permitted quantitative 3D mapping at a high and near-isotropic spatial resolution (2.3/2.0 μm laterally/axially). The dual-view geometry was essential to obtain accurate and reliable tissue/cavity volume measurements across the whole animal. Figure S4A shows the schematic diagram of the DV-OCM system, which incorporates two identical illumination/detection arms that are symmetric with respect to the intermediate plane between the two objective lenses. The collimated illumination was first focused by a scanning lens (*f* = 50 mm, Plossl lens build from two *f* = 100 mm achromats, EdmundOptics, 47-317). Then a knife-edge right angle prism mirror (Thorlabs, MRAK25-P01) was used to equally split the probe beams into the two scanning arms. Each optical arm contained a tube lens (Thorlabs, TTL200-S8) and an objective lens (Nikon, Plan Fluor 4x, NA 0.13) both of which were carefully co-aligned to focus the light on the same spot on the sample plane. The laser source for OCM imaging was a supercontinuum laser (YSL Supercontinuum Source SC-PRO 7), of which the spectrum was custom-filtered to cover the band 760-920nm (center wavelength 832 nm), and fed into a 20/80 fiber-coupler based interferometer. A custom-built k-wave spectrometer (Lan and Li, 2017) recorded the interference signals on a 2048 pixels, 28 kHz line camera (AViiVA SM2 CL, e2v, Cedex, France). The measured lateral and axial resolution in tissue (n=1.35) was ~2.3 and 2.0μm, respectively. A profile of depth scan (A-line) was reconstructed from the interference spectral signal following a regular OCT postprocessing procedure including dispersion compensation, background subtraction, spectrum reshaping and inverse fast Fourier transformation (Liba et al., 2016) and was saved as individual TIFF files using a custom Matlab (v2019b) script. Each cross-sectional image (B-scan) contained 512 A-lines and was split into two images obtained from the different views. In order to record a single 3D image stack, 256 B-scans were performed which took ~ 4.7s in total. The image acquisition was controlled by a custom-written Labview program which synchronized the galvo scanners (Cambridge, VM500+) and read-out the spectrometer via an FPGA card (NI, NI PCIe-7841R).

For imaging, live non-anesthetized larvae and polyps were sandwiched between two coverslips with spacers and mounted on a stage with manual x,y,z movement.

The raw OCM images were post-processed using FIJI (Schindelin et al., 2012) (v1.53c) by median filtering with 2 pixel width to remove speckle noise followed by a re-scale operation to correct for the difference between the axial and lateral voxel dimensions. Alignment was performed using Fyama (automatic registration, block matching, default settings) (Fernandez and Moisy, 2020). The two views were combined into one volume and then segmented into tissue and cavity using the trainable WEKA segmentation FIJI plugin (Arganda-Carreras et al., 2017). Tissue volume and cavity were calculated from the segmented masks using the IntrinsicVolumes3D function from the MorpholibJ package (Legland et al., 2016). 3D views were created with VolumeViewer (2.01) (Barthel, 2012) in FIJI (https://imagej.nih.gov/ij/plugins/volume-viewer.html).

### Pressure measurements

The 900A micropressure system (World Precision Instruments, SYS-900A) was used to perform direct measurements of body cavity pressure according to the manufacturer’s instructions and adapted from previous work (Chan et al., 2019; Petrie et al., 2014). A pre-pulled micropipette of tip diameter 1μm (WPI, TIP1TW1) was filled with 1M KCl solution using a MicroFil flexible needle (WPI, MF34G-5), placed in a microelectrode holder half-cell (WPI, MEH6SF) and connected to a pressure source regulated by the 900A system (WPI, 900APP). Prior to making a pressure measurement, each new microelectrode was calibrated using a calibration chamber (WPI, CAL900A) filled with 0.1M KCl solution. The microelectrode and a reference electrode (WPI, DRIREF-2) were then mounted onto the micromanipulator (Narishige MO-202D) within an inverted Zeiss Axio Observer microscope.

To prepare the animals for pressure measurement, polyps were first placed onto 6-cm Petri dishes in ASW with or without pharmacological inhibitors, and allowed to adhere to the bottom surface. The dish was mounted on the microscope with the reference electrode immersed in the medium, and the microelectrode was then lowered into the sample dish and inserted into the cavity of the animal by piercing through the muscular body wall. The microelectrode was then maintained in place to record the pressure reading for up to 8-10 minutes, ensuring that the tip was visible. A time-lapse video of the animal was recorded simultaneously with the measurement of pressure to capture contractions. Data that showed a very rapid increase or decrease in pressure within 10s of probe insertion were discarded, as these indicate blockage of the micropipette during insertion or substantial leakage through rupture, respectively. The change in cavity pressure was then plotted as a function of time, overlaid with the time-lapse video to identify the period of muscle contractions.

### Pharmacological treatments

Budded larvae were treated with 5 mM hydroxyurea (HU) (Sigma, h8627) in ASW while controls were incubated in 12 ppt ASW. For high-throughput imaging, animals were transferred in a 384 well-plate containing 25 μl 5 mM HU or 25 μl 12 ppt ASW for controls. Animals were imaged at 5-minute time resolution for up to 80 hours at 27 °C using the Acquifer microscope with the brightfield channel. For immunostaining, animals were kept in glass dishes at 27 °C and were fixed after 3 and 24 hours of incubation.

To inhibit muscle-driven body contractions, animals were incubated with one of the following drugs: 1 mM linalool (Goel et al., 2019) (Sigma, L2602) with 0.03% DMSO, 400 μM menthol (Abrams et al., 2015; Batham and Pantin, 1950b) (Sigma, M2772), with 0.1% DMSO, 0.5% MgCl_2_, or 0.5 mM rocuronium bromide (Khuenl-Brady and Sparr, 1996) with 0.2% DMSO (Sigma, R5155) in glass dishes for three days. As a control, animals were incubated in either 12 ppt ASW or 12 ppt ASW with 0.2% DMSO. For the BMP KD experiment in combination with inhibition of muscle contraction, we injected fertilized eggs with BMP shRNA (550 ng/μl). Animals were either fixed after 2 days (t0) or incubated in 0.1% DMSO (control), 400 μM menthol with 0.1% DMSO, or 1 mM linalool with 0.03% DMSO for 4 days. All dishes were sealed with parafilm without refreshing the drug solutions. Fixed samples were stained with phalloidin Alexa Fluor 546 (Thermo Fisher, A22283, 1:100), and mounted in glycerol on microscope slides. All samples were imaged using a Zeiss LSM 780 NLO or LSM 880 confocal inverted microscope with Plan-Apochromat 20x/0.8 objective. Body length was measured by drawing a segmented line from the oral to the aboral pole in the midplane of the animal. Area of body cavity and internal tissues in the mid-plane section was measured in FIJI by manual segmentation.

### Fixation and immunostaining

Animals were anesthetized in 7% MgCl_2_ prior to fixation. Fixation was performed for 1 hour at room temperature either with 4% paraformaldehyde (EMS, E15710) in PBS with 0.1% Tween (PTw 0.1%, Sigma, P1379) for phosphorylated histone 3 antibody staining (Chen et al., 2020) (pH3, Sigma # 05-806, 1:100), or with Lavdovsky’s fixative (3.7% formaldehyde, 50% ethanol, 4% acetic acid) for Cadherin3 antibody detection (gift from Technau lab (Pukhlyakova et al., 2019), 1:500). Animals were permeabilized in 10% DMSO (Thermo Fisher, 85190) in PBS for 20 minutes and washed with PBS with 0.2% Triton (Sigma, T8787) (PTx 0.2%), followed by 1-hour incubation in blocking buffer containing PTx 0.1%, 0.1% DMSO, 1% BSA (Sigma, A2153), and 5% Goat serum (Sigma, G9023). Samples were incubated with the primary antibody in blocking solution overnight at 4 °C. After washing with PTw 0.1%, animals were incubated with the secondary antibody goat-anti-mouse Alexa-488 (Thermo Fisher, A-11001, 1:500) or donkey-anti-mouse Alexa-488 (Thermo Fisher A21202, 1:500) in PTx 0.1% overnight at 4 °C. For (additional) staining for F-actin and nuclei, phalloidin Alexa Fluor 546 (Thermo Fisher, A22283, 1:100) and Hoechst 34580 (Sigma, 63493, 1:1000) were used, in PTx 0.1% overnight at 4 °C. Finally, animals were washed in PTw 0.1% and cleared in 80% glycerol (Merck).

### Confocal imaging

Samples were imaged using a Zeiss LSM 780 confocal inverted microscope with Plan-Apochromat 20x/0.8 objective, or using a Zeiss LSM 880 point scanning confocal microscope controlled with the Zeiss Zen 2.3 (black edition) software, with Plan-Apochromat 20x/0.8 air objective. To image epidermal cells, we used a Plan-Apochromat 40x/1.4 Oil DIC objective, AiryFast mode and tile scans, at 0.5 μm z-resolution. Depending on the staining, we used laser lines diode 405 nm, argon multi-line 458/488/514 nm and/or HeNe 561 nm.

### Quantification of cellular properties

The number of pH3 positive-mitotic cells was counted manually in FIJI and normalized to the imaged tissue volume. Epidermal thickness was measured manually in images of the mid-plane of the animal at multiple locations along the body axis using FIJI. The measurement was taken in the mid-body region between 30% and 70% of the total length of the oral-aboral axis. Quantifications of the cell apical surface area and angle with respect to the body axis were based on the images taken with the 40x magnification objective. To correct for the typical uneven surface of the animals and the body column deflation upon addition of glycerol, the Minimum cost Z surface projection FIJI plugin (Li et al., 2006; Lombardot, 2017, available at https://imagej.net/Minimum_Cost_Z_surface_Projection) was used to obtain separate, flattened layers for the ectoderm tissue and the underlying muscular tissue. Next, one or more flattened slices showing cell boundaries were selected and used as input for detection of cells by Cellpose (Stringer et al., 2021) (2D mode, model-type cytoplasm). The mask produced by Cellpose was converted to a label image and was manually corrected. For phalloidin staining in polyps, correctly segmented cells were manually selected. Cell area, ellipse elongation and orientation were obtained using the MorphoLibJ package (Legland et al., 2016). To calculate the cell angles with respect to the body axis, a line was drawn from the oral to the aboral pole along the midline of the animal. Angles for each location on this line were calculated for 10-pixel segments. For each cell, the nearest position on the line was determined, and the angle of the oral-aboral line at this point was used as a reference angle to compare to the cell’s angle.

### shRNA design and synthesis

shRNAs were designed based on (He et al., 2018) and using the siRNA Wizard from Invivogen (available at https://www.invivogen.com/sirnawizard/design_advanced.php). Primers were synthesized by Sigma and IDT.

Primers were annealed at 98 °C for 5 min in the PCR machine or heat block and allowed to cool down to room temperature. shRNA was synthesized using the T7 MegaShortScript kit (Invitrogen, AM1354) with an incubation time of 6 hours, followed by a purification step using magnetic SPRISelect beads (Beckman Coulter B23319) in the presence of 46% isopropanol (Fishman et al., 2018). Samples were incubated for 15 minutes at room temperature, and were then placed in magnetic stands for 5 minutes, until the solution appeared clear. The samples were washed twice using 80% freshly prepared ethanol, after which they were shortly dried and resuspended in RNase free water. The solution was aliquoted and stored at −80 °C.

### Microinjection of shRNAs

Unfertilized eggs were dejellied in 4% cysteine solution (Sigma, 168149) in ASW for 9 minutes and washed with ASW. Eggs were then fertilized and injected with shRNA targeting GFP, Tbx20 or BMP (500-1500 ng/μl), combined with Texas Red-labelled Dextran (ThermoFisher, D3328) using a Femtojet Express (Eppendorf). Injected eggs were kept at room temperature and transferred to 27 °C the following day.

### CRISPR/Cas9

Single guide RNAs (sgRNAs) were designed using the online web interface http://chopchop.cbu.uib.no (Labun et al., 2019). Two gRNAs targeting the coding sequence of both Tbx20 paralogs were commercially synthesized (Sigma aldrich). Recombinant Cas9 protein (900 ng μl−1; PNA Bio, #CP01-20) was co-injected with both sgRNAs (20μM each) into unfertilized Nematostella oocytes. Injected oocytes were then fertilized and raised at room temperature for imaging and sequencing.

### Photoconversion of Kaede

mRNA was synthesized using the HiScribe™ T7 ARCA mRNA Kit (with tailing) (NEB, E2060S) and a PCR product amplified from the Kaede-NLS plasmid (addgene, 57319) (primers are shown in key resources table). Following mRNA purification with magnetic SPRISelect beads (Beckman Coulter B23319), fertilized eggs were co-injected with a solution mix containing Kaede-NLS mRNA (200 ng/μl), shRNA targeting BMP (430 ng/μl) or Tbx20 (560 ng/μl), and FITC (ThermoFisher, 46425). Control embryos were only injected with Kaede-NLS mRNA and FITC. Photoconversion of Kaede (Ando et al., 2002) was performed on anesthetized larvae on a Zeiss LSM 780 microscope with Plan-Apochromat 20x/0.8 objective, using the ‘bleaching’ and ‘regions’ option, and using the 405 laser at 0.6% laser power, with 80 iterations. Imaging of developed animals after photoconversion was performed on a Zeiss LSM 780 or 880, with Plan-Apochromat 20x/0.8 objective.

### Biophysical manipulation

Glass capillaries (TW100F-4, World Precision Instruments) were pulled using a micropipette puller to obtain approximately 10μm diameter tubes. To create openings across the tube length, pulled capillaries were mounted in an imaging slide with double-side tape, and laser-cut holes were created using the 355 nm laser of the Olympus FV1200 microscope. During the piercing of the larvae, open and closed capillaries were handled with a joystick for micro-injection (MO-202U, Narishige-group). For open capillaries, the pressure was equalized using the compensation pressure function of the Eppendorf electronic microinjector, the FemtoJet®.

### Data analysis

Data analysis and generation of plots was performed in RStudio version 1.1.442 (R Core Team, 2019; RStudio Team, 2015), making use of the tidyverse (Wickham et al., 2019), ggplot2 (Wickham, 2009), ggpubr (Kassambara, 2020), gtable (Wickham and Pedersen, 2019), and ggrastr (Petukhov et al., 2020) packages and Prism 8. Wilcoxon rank sum tests were used for significance testing, unless mentioned otherwise. The forecast package (Hyndman et al., 2008, 2020) was used to remove outliers in time sequence data and the zoo package (Zeileis and Grothendieck, 2005) was used to calculate the rolling sum in time series data.

### Biophysical model

In the limit of small strains, the axial and azimuthal strains can be written as: 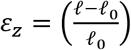 and 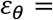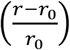, respectively, so that the passive stresses in the axial and azimuthal directions read as 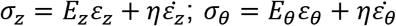 where *E*_*z*_ is the axial elastic modulus and *E*_*θ*_ is the azimuthal elastic modulus of the wall, and *η* is the effective viscosity of the wall (assumed to be isotropic). All asymmetries in the relative stiffnesses in the principal directions can be ascribed to the difference in the densities of the azimuthal muscle fibers versus the axial muscle fibers. For simplicity, we assume that the passive elastic moduli are of the same order as the active moduli with *E*_*z*_ ≈ *E*_*mz*_ and *E*_*θ*_ ≈ *E*_*mθ*_ so that the active muscular stresses in the axial and azimuthal direction can be modeled as σ_*mz*_ = *E*_*mz*_*ε*_*z*_, σ_*mθ*_ = *E*_*mθ*_*ε*_*θ*_ (McMahon, 2020). Then the axial strain rate 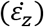 and azimuthal strain rate 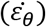 for the tissue are given by:

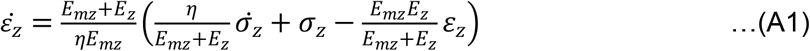

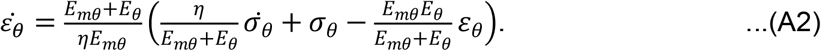

For the simplest case, we assume that the volume of the wall tissue (*V*_*w*_) as well as the cavity volume (*V*_*c*_) remain constant during the shape evolution of the larva to polyp,

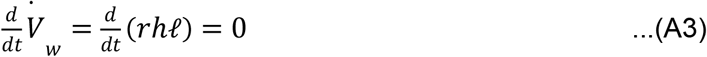

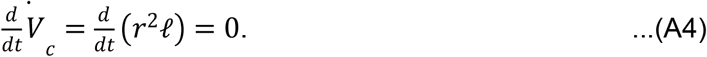

In this limit, Eqs. A3–4 imply that as the radius decreases, the length of polyp can increase with negligible changes in the wall thickness. The final shape of the polyp is obtained from solving Eqs. (A1–4) with the boundary conditions *ℓ*(0) = *ℓ*_0_, *r*(0) = *r*_0_, and *h*(0) = *h*_0_, and typical parameter values (Alexander, 1962):*E*_*θ*_ = 1000 *Pa*; *E*_*mθ*_ = 10 *Pa*; *E*_*z*_ = 10 *Pa*; *E*_*mz*_ = 1 *Pa*; *η* = 10^4^ *Pa*.*s*; *p*_0_ = 0.1 *Pa*; *p*_*T*_ = 5 *Pa*; *ω* = 1 *Hz*; *r*_0_ = 100*μm*; *ℓ*_0_ = 500*μm*; *h*_0_ = 10*μm*, and yields the results shown in Figure 6A and 6B.

At steady state, the strain rates in Eq. (A1-2) vanish, and the body wall tissue volume remains conserved which implies that *rhℓ* = *const*. In Tbx20 KD animals, we expect that the elastic stiffness resulting from the disorganized parietal and circular muscles becomes negligible, so that the axial and azimuthal stiffnesses are similar, i.e, *E*_*z*_ ≈ *E*_*θ*_. With these assumptions, Eqs. A1–2 yields *ε*_*θ*_ = *ε*_*z*_, i.e. Tbx20 KD larvae will isotropically inflate via fluid uptake to form a spherical polyp morphology. In contrast, BMP KD animals only maintain the circular muscles, implying that effective stiffness in the azimuthal direction is greater than the stiffness in axial direction, *E*_*θ*_ ≫ *E*_*z*_. Then the equilibrium relationships (A1-2) simplify to: *E*_*θ*_*ε*_*θ*_ = 2*E*_*z*_*ε*_*z*_, and furthermore both cavity and wall tissue volumes conserved, i.e. *rhℓ* = *const* and *r*^2^*ℓ* = *const*. These simple relations explain the origin of a thin and elongated morphology with a conserved cavity volume (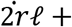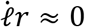), and large aspect ratio 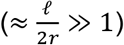, while having negligible changes in body wall thickness. Finally, wild type animals inflate their cavity through fluid uptake 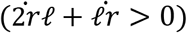, while the body wall tissue volume does not change (*rhℓ* = *const*). As a consequence, a similar situation to the BMP KD animals is obtained, i.e. the aspect ratio 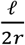 can be larger than unity if the axial stiffness is lower than the azimuthal stiffness, i.e *E*_*z*_/*E*_*θ*_<1, since then the body geometry will imply that the axial strain will be larger than the azimuthal strain, i.e.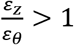.

### Physical balloon experiments

For the physical simulacrum experiments, we used latex balloons (12” Balloons, red, Celebrate It™, Item No: 10108443), elastic braided bands (1/4” Braided Elastic Hank, Loops & Threads™, Item No: 10187887), inextensible electrical tapes (3/4”, pressure sensitive vinyl tape, red, Gardner Bender) and stretchable adhesive (Loctite vinyl fabric & plastic flexible adhesive, Item No: 1360694). To mimic the various Nematostella phenotypes, our initial starting point is identical, the uninflated latex balloon (Figure S10B), which when inflated using compressed air, expands isotropically with aspect ratios 1, serving as a control for the following mimic experiments (Figure S10B). To obtain the Tbx20 KD phenotype, we first inflate the balloon such that it extends basally (volume ~ V_0_). We then stick small pieces of extensible tapes in randomized orientation (here, tapes serve as proxy for muscles on the elastic wall of the balloons), following further inflation (volume ~ V_f_), thereby obtaining close to spherical shapes (aspect ratios in the range of *ℓ*/d~1.1- 1.3 and volume change ratios in the range of V_f_/V_0_ ~2.5-5.5) (Figure S10C). Wildtype mimics are obtained using two methods: (I) the balloon is inflated basally, followed by attaching thin elastic bands azimuthally with the stretchable adhesive. The elastic bands make the azimuthal stiffness of the balloon larger than its axial stiffness. When this balloon is inflated, it elongates longitudinally, mimicking the shape changes in wild type polyps with aspect ratios *ℓ*/d~2 and volume changes V_f_/V_0_~2-4 (Figure S10D). The second method (II) to obtain wildtype mimics employs a slightly different approach wherein the balloon is inflated more at the beginning, sealed, and then stretched longitudinally while attaching inextensible tapes helically. This determines the volume change at the beginning and is followed by remodeling of the balloon wall to yield slender shapes (*ℓ*/d~4.2-5 and V_f_/V_0_~2.5-4, here V_f_/V_0_ is obtained by comparing the inflated balloon stage before remodeling with the basally inflated stage) (Figure S10E). To obtain the BMP KD mimic, we needed to obtain much larger aspect ratios, and it is known from experiments that the cavity volume remains constant during the transformation of polyps. We first inflated the balloons to a basally distended shape and sealed the opening of the balloon. The inflated balloon mimicking the larval stage was then stretched axially and taped with inextensible pressure sensitive vinyl tapes (electric tapes) to constrain the azimuthal direction while elongating in the axial direction. This led to high aspect ratios *ℓ*/d~7.5 while keeping the cavity volume constant (V_f_/V_0_=1) (Figure S10F). Using the simple toolbox of balloons, elastic bands, electrical tapes and adhesives, we are able to recapture the steady state phenotypes and the phase space of *Nematostella* animals.

## Notes

### Competing Interest Statement

The authors have declared no competing interest.

### Summary of Updates

In the abstract, "habits" should be "habitats".

