## Supplemental information for "Ethology of morphogenesis reveals the design principles of cnidarian size and shape development"

### Supplemental figures

### Supplemental videos

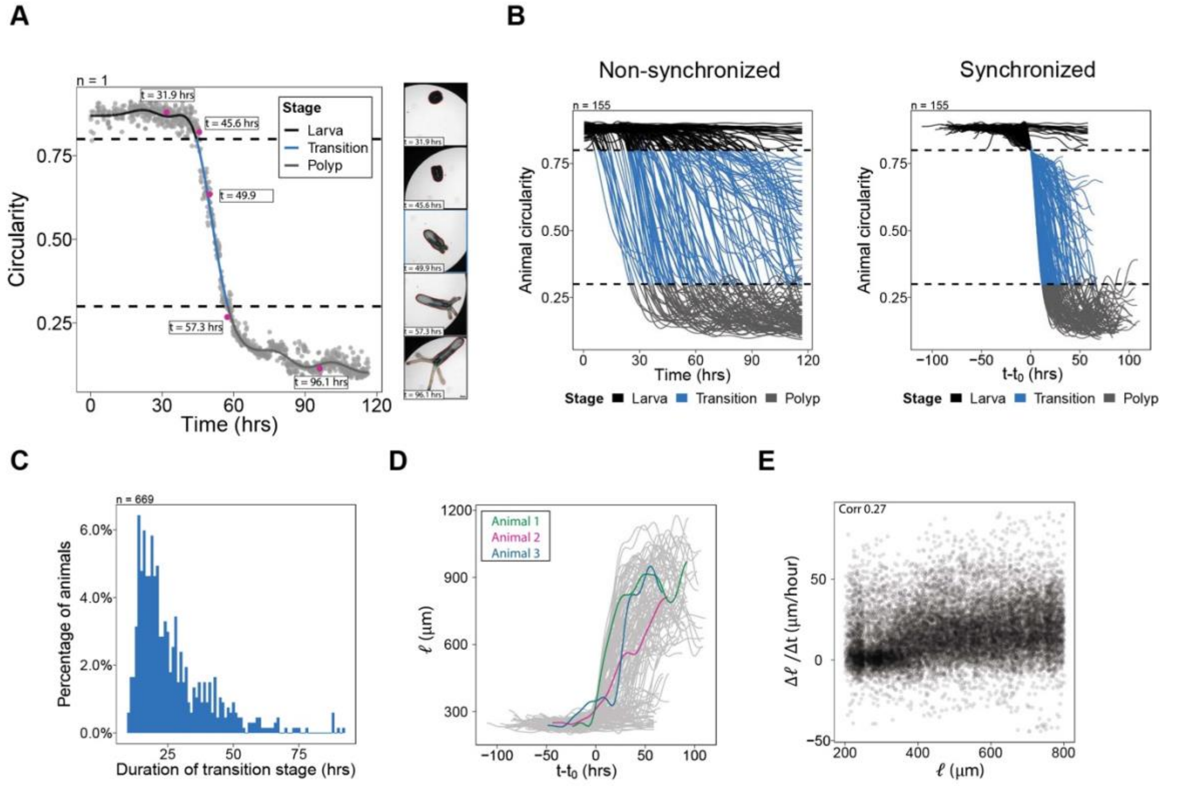


**Figure S1. Developmental staging based on the changes of organism circularity.**
(A) Left, raw measurements (dots) and smoothed curve of animal circularity over time. A threshold set at 0.8 and at 0.3 marks distinct stages in development (larva, transition, polyp). Right, brightfield snapshots for different time points (pink dots).
(B) Left, smoothed trajectories for circularity over time in the same experiment (*n = 155* animals). Right, smoothed trajectories for circularity, shifted in time to synchronize the start of the transition stage. $t_0$ marks the start of the transition stage.
(C) Histogram for the duration of the transition stage, obtained from combined data from 5 different experiments (*n = 669* animals total).
(D) Smoothed curves for time-shifted body length, highlighting three individuals with different elongation dynamics.
(E) Weak correlation between body length and change in body length (pearson correlation coefficient).

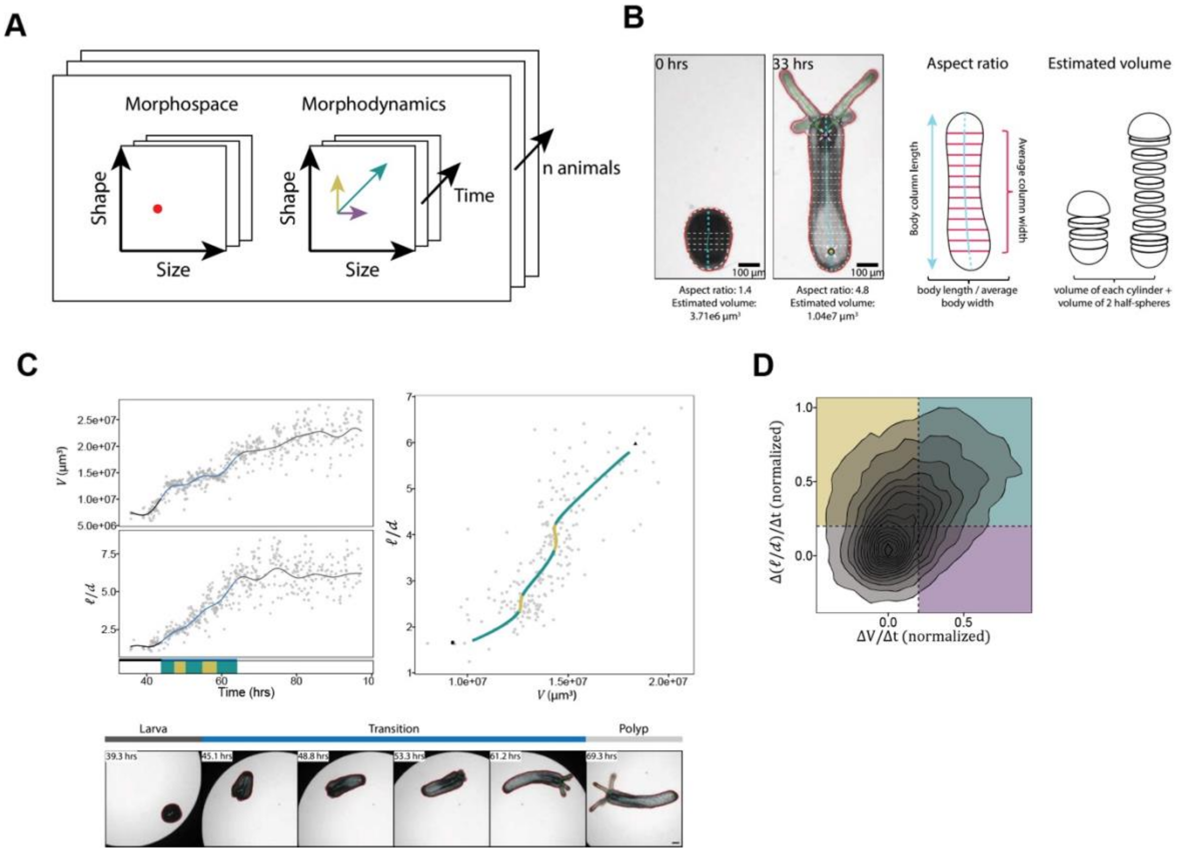


**Figure S2. Quantifications of morphospace and morphodynamics.**
(A) Schematic showing morphospace defined by the dimensions shape and size, and morphodynamics defined by the changes in shape and size. Time and individual animals are additional dimensions.
(B) Left, brightfield images of a larva and the corresponding polyp, with overlaid annotations. Right, schematic showing how aspect ratio and body column volume are estimated from 2D brightfield images.
(C) Example of one individual animal showing measurements in time and in morphospace. Colors indicate stage and morphodynamics (yellow: organismal convergent extension, purple: isotropic expansion, green: anisotropic expansion, grey: no elongation). Snapshots are shown below.
(D) Density plot of morphodynamics in the transition stage. Peak level corresponds to a probability of 0.125, each contour line indicates 0.83% probability.
Scale bars: 100 µm (B and C).

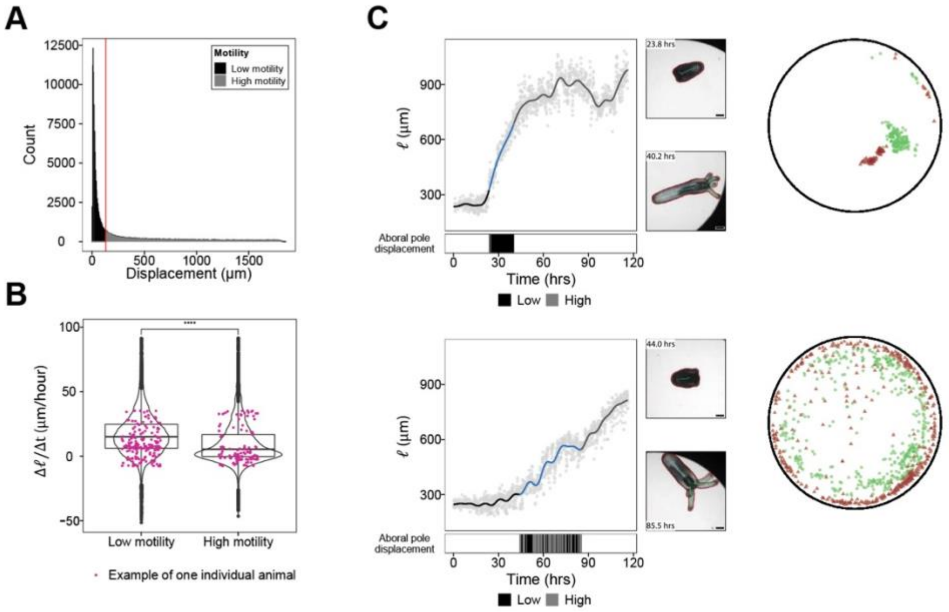


**Figure S3. Elongation dynamics in sessile and motile larvae.**

(A) Histogram of measured displacement per 5-minute interval for all animals combined. A displacement of up to 130 µm per 5 minute-interval is considered low (black), more than 130 µm is considered high (light gray).

(B) Violin plot showing the distributions for body length change for time points with low or high motility, respectively (two-sided unpaired Wilcoxon rank sum test; ****$p < 0.0001$). Animals can switch between high and low motility, and indicated by the pink dots, corresponding to one individual.

(C) Left, measurements for body length over time for a sessile (top) and motile (bottom) animal, showing different elongation dynamics. Motility is given by aboral pole displacement (bars). Snapshots before and after elongation are shown in the middle. Right, projection of oral and aboral pole locations in the well during the transition stage.

Scale bars: 100 µm.

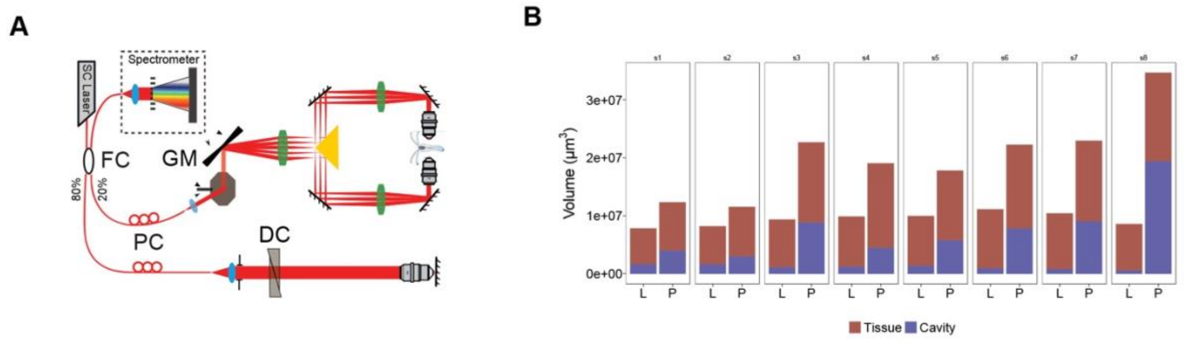


**Figure S4. Measurements of volumetric changes in the larva-polyp transition.**
(A) Schematic diagram of dual-side view spectral-domain Optical Coherence Microscopy. FC: fiber coupler; GM: galvo mirror; DC: dispersion compensator; PC: polarization controller; SC Laser: supercontinuum laser.
(B) OCM-based measurements of tissue (red) and cavity volume (blue) for 8 individuals, before and after larva-polyp transition.

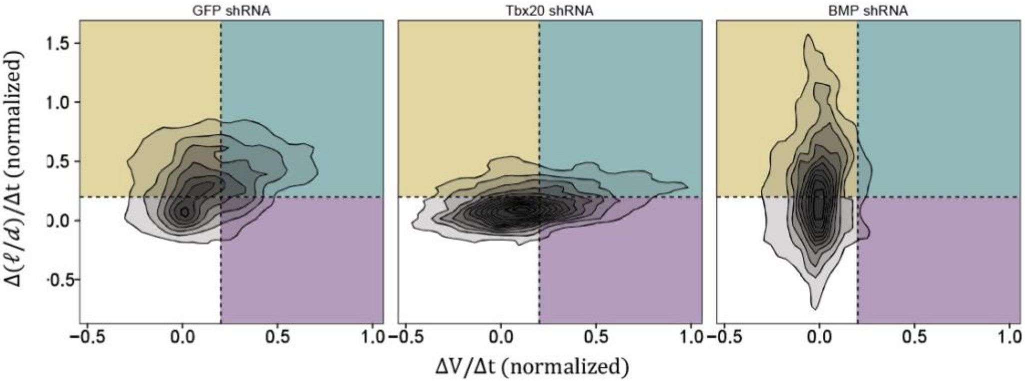


**Figure S5. 2D Density plot of morphodynamics for control, Tbx20 KD and BMP KD animals.** Peak density corresponds to 0.14, each contour line indicates a probability of 1.1%. Color-coded for morphodynamics: organismal convergent extension (yellow), isotropic expansion (purple), anisotropic expansion (green), no elongation (grey) (control *n = 74* animals, Tbx20 KD *n = 40* animals and BMP KD *n = 40* animals). Scores are calculated by normalizing to the 95% percentile values.

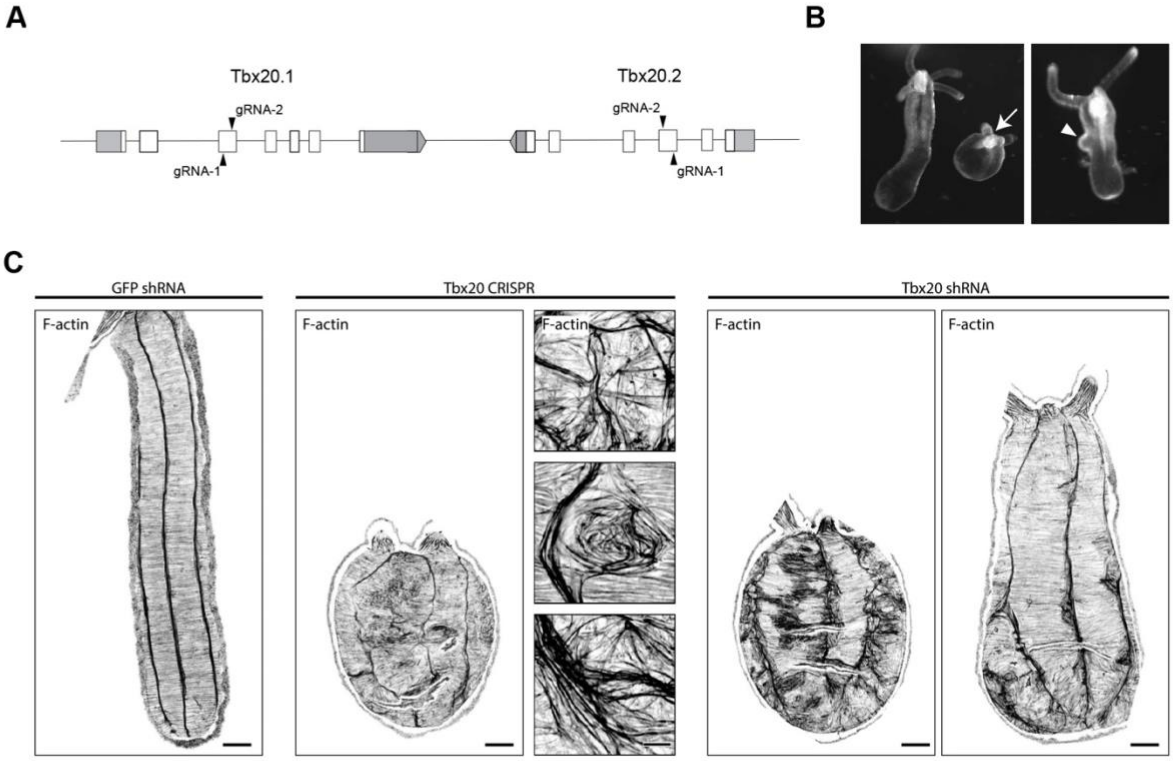


**Figure S6. Generation and characterization of Tbx20 loss of function.**
(A) Schematic showing the gRNA design to generate F0 Tbx20 CRISPR mutants.
(B) Examples of Tbx20 CRISPR/cas9-injected animals showing round morphologies (white arrow) and a bleb-like structure (white arrowhead).
(C) Comparison of muscle organization in a control animal, Tbx20 CRISPR/cas9-injected animals, and Tbx20 shRNA KD animals. Images show maximum intensity projections of stacks that are straightened using the MinCostZProjection plugin in FIJI.
Scale bars: 50 µm (C, overview images) and 10 µm (C, zoom-ins for Tbx20 CRISPR/cas9-injected animals).

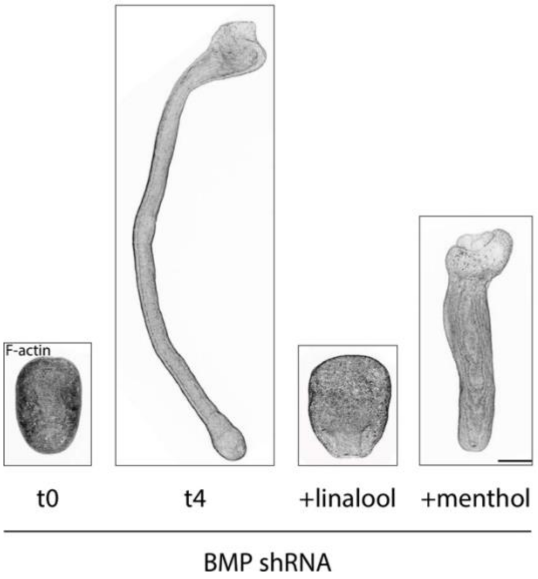


**Figure S7. Pharmacological inhibition of muscle function in BMP shRNA KD animals prevents body elongation. (A)** Maximum intensity projections of confocal images of BMP shRNA injected animals fixed and stained with phalloidin at 2 dpf (t0), or 6 dpf (t4, drug-treated and control). Quantification is shown in Figure 4E.
Scale bar: 100 µm.

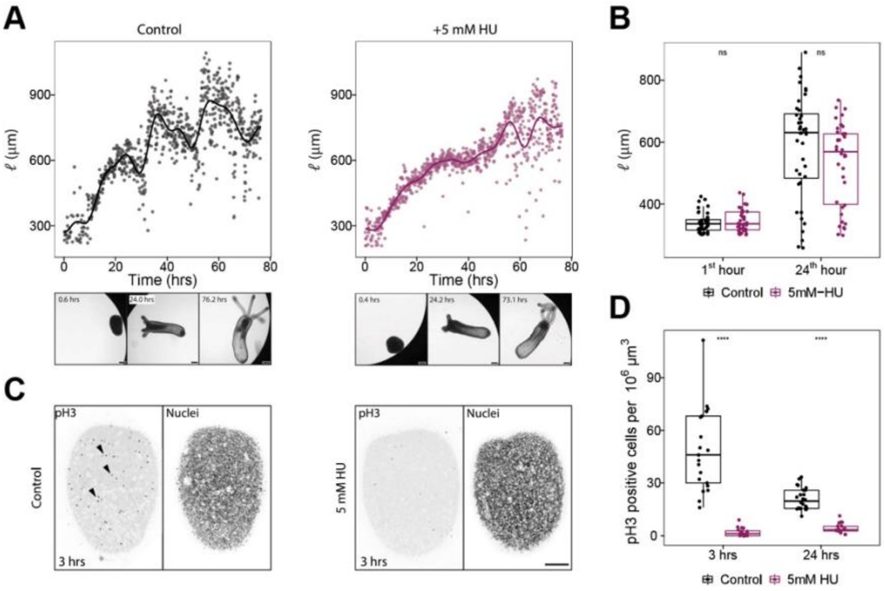


**Figure S8. Inhibition of cell proliferation in budded larvae does not prevent axial elongation.**
(A) Measurements for body column length for a control animal (left) and an animal treated with 5 mM hydroxy urea (right). Illustrative bright field images are shown below the plots for 3 time points.
(B) Quantification of body length after one hour and 24 hours since the start of treatment, measured in budded larvae (two-sided unpaired Wilcoxon rank sum test; n.s $p > 0.05$) (control: $n = 42$, 5mM HU: $n = 42$).
(C) Maximum intensity projection of a control larva (left) and a larva treated with 5 mM HU for 3 hrs, stained for pH3 and for nuclei (hoechst). Arrowheads mark examples of pH3 positive nuclei. (D) Quantification of the number of pH3 positive cells per $10^6$ µm$^3$ (two-sided unpaired Wilcoxon rank sum test; ****$p < 0.0001$) (control 3 hrs: $n = 21$, 5mM HU 3 hrs: $n = 19$, control 24 hrs: $n = 24$, 5 mM HU 24 hrs: $n = 20$).
Scale bars: 100 µm (A), 50 µm (C).

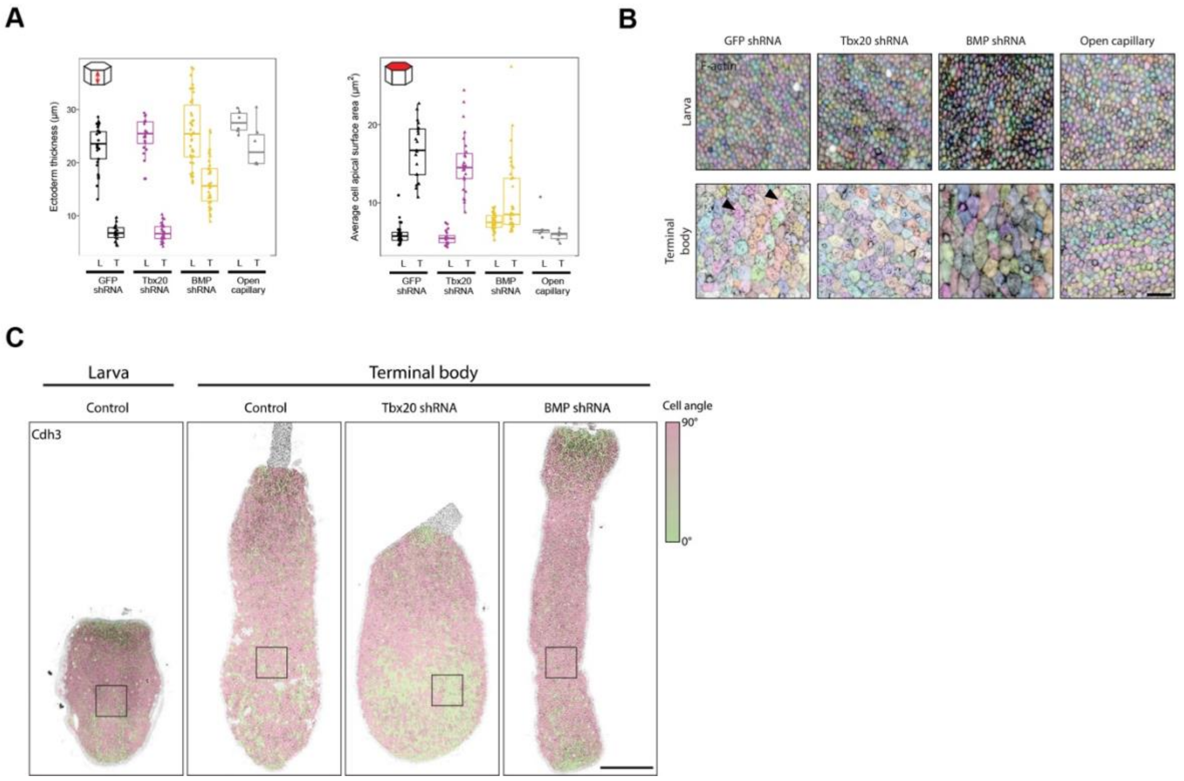


**Figure S9. Cellular behaviors under distinct muscular hydraulic stresses.**
(A) quantification of epidermal thickness (left) and average apical cell surface area (right) for larva (L, control *n = 29*, Tbx20 KD *n = 20*, BMP KD *n = 38*, and open capillary *n = 6*) and terminal body (T, control *n = 21*, Tbx20 KD *n = 27*, BMP KD *n = 34*, and open capillary *n = 6*).
(B) Epidermal apical surface view of animals stained for F-actin. Note the actin-enriched rings in the apical surface area of epidermal cells (black arrowheads) in wild type and Tbx20 KD animals. This feature of polyp cell differenciation is lacking in animals pierced with an open capillary. Images are flattened using the MinCostZProjection FIJI plugin. Colors indicate cell segmentation.
(C) Epidermal surface view for animals stained for Cdh3. Colors indicate the cell angle with respect to the oral-aboral axis (magenta = aligned with body axis, green = perpendicular to the body axis).
Scale bars: 10 µm (B), 100 µm (C).

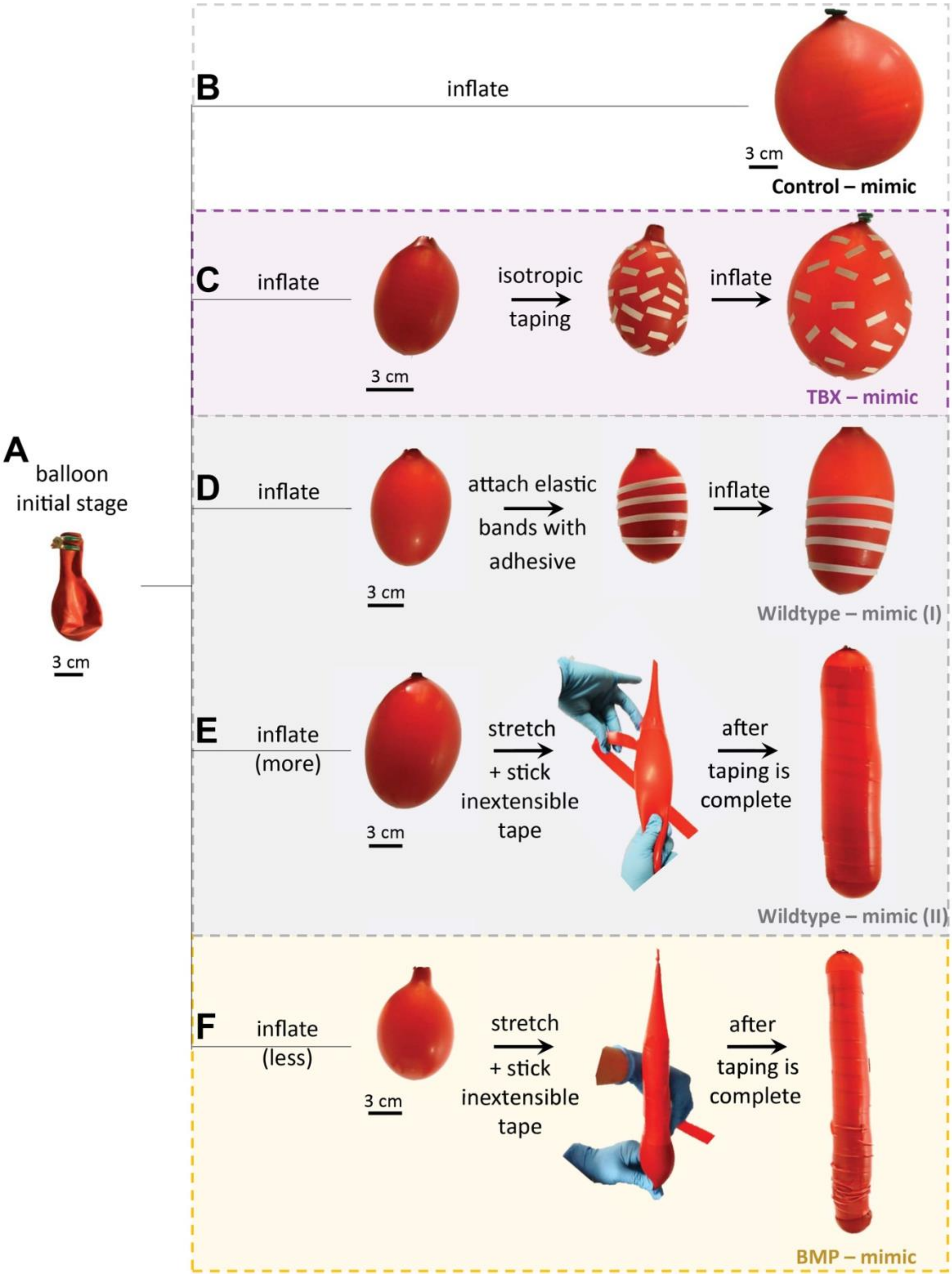


**Figure S10. Physical balloon model experiments mimic the phenotypes observed in *Nematostella* control and shRNA KD animals**.
(A) A latex balloon at its initial state before inflation, which is the starting point for all our simulacrum experiments.

(B) When it is inflated using compressed air, the balloon expands isotropically (aspect ratio, ℓ/d~1), representing the control for the physical simulacrum mimic models for *Nematostella* phenotypes.
(C) To generate a Tbx20 KD mimic, we first inflate the balloon such that it extends basally. We then stick small pieces of extensible tapes in randomized orientation (here, tapes serve as proxy for muscles on the elastic wall of the balloons), followed by further inflation to obtain a spherical shape mimicking the Tbx20 KD phenotypes (ℓ/d~1.25).
(D-E) Wildtype mimics are obtained using two methods: (D) (I) the balloon is inflated basally, followed by attaching thin elastic bands azimuthally with a stretchable adhesive and then inflating it further. The elastic bands make the azimuthal stiffness larger than the axial direction, thereby allowing expansion of the balloon longitudinally. This obtains both slender shapes as well as volume expansion for the wildtype mimic (I) (ℓ/d~2). (E) (II) the balloon is inflated more than in (D) followed by stretching it longitudinally and helically attaching inextensible tapes (II) (ℓ/d~4.3).
(F) To obtain the BMP KD mimic, we used the same protocol as in (E) but start with a low volume balloon. We stretch and tape the balloon helically. The low initial volume allows for much larger aspect ratios for the final BMP KD mimics (ℓ/d~7.5).

### Supplemental video titles

**Video S1.** Brightfield videos of a selection of developing *Nematostella* larvae from the same experiment.

**Video S2.** Brightfield recording with image annotations and quantification of body length over time for a sessile animal.

**Video S3.** Brightfield recording with image annotations and quantification of body length over time for a motile animal.

**Video S4.** Brightfield recording with image annotations and quantification of animal volume showing muscle driven pumping behavior and cavity inflation.

**Video S5.** Brightfield recording with image annotations and quantification of aspect ratio over time in primary polyps before and after treatment with 1.1 mM linalool.

**Video S6.** Brightfield recording of a developing Tbx20 KD animal.

**Video S7.** Brightfield recording of a developing BMP KD animal.

**Video S8.** Brightfield recording and quantification of cavity pressure over time for a contracting polyp.

**Video S9.** Brightfield recording and quantification of cavity pressure over time for an animal treated with 7% $MgCl_2$.
